# Identification and characterization of key long non-coding RNAs in the mouse cochlea

**DOI:** 10.1101/2020.07.10.197251

**Authors:** Tal Koffler-Brill, Shahar Taiber, Alejandro Anaya, Mor Bordeynik-Cohen, Einat Rosen, Likhitha Kolla, Naama Messika-Gold, Ran Elkon, Matthew W. Kelley, Igor Ulitsky, Karen B. Avraham

## Abstract

The auditory system is a complex sensory network with an orchestrated multilayer regulatory program governing its development and maintenance. Accumulating evidence has implicated long non-coding RNAs (lncRNAs) as important regulators in numerous systems, as well as in pathological pathways. However, their function in the auditory system has yet to be explored. Using a set of specific criteria, we selected four lncRNAs expressed in the mouse cochlea, which are conserved in the human transcriptome and are relevant for inner ear function. Bioinformatic characterization demonstrated a lack of coding potential and an absence of evolutionary conservation that represent properties commonly shared by their class members. RNAscope analysis of the spatial and temporal expression profiles revealed specific localization to inner ear cells. Sub-cellular localization analysis presented a distinct pattern for each lncRNA and mouse tissue expression evaluation displayed a large variability in terms of level and location. Our findings establish the expression of specific lncRNAs in different cell types of the auditory system and present a potential pathway by which the lncRNA *Gas5* acts in the inner ear. Studying lncRNAs and deciphering their functions may deepen our knowledge of inner ear physiology and morphology and may reveal the basis of as yet unresolved genetic hearing loss-related pathologies. Moreover, our experimental design may be employed as a reference for studying other inner ear-related lncRNAs, as well as lncRNAs expressed in other sensory systems.

## Introduction

Hearing loss, or deafness, is a heterogeneous pathology caused by congenital or acquired factors, and affecting an estimated 466 million people worldwide [1]. Hearing loss can have significant implications on communication, quality of life and educational attainments [2], highlighting the importance of early diagnosis toward optimizing patient management and therapeutic outcomes [3]. The auditory system and the hearing process are complex, with multiple genes and pathways involved in their development and maintenance [4]. Nearly all variants identified as responsible for hearing loss are located in the coding regions of genes, although many more regulatory mutations probably exist in non-coding regions than have currently been reported [5–7]. Since hearing is very likely to be significantly influenced by regulatory regions of the genome, identifying these elements is of critical importance.

Long non-coding RNAs (lncRNAs) are transcripts greater than 200 bp in length that do not produce functional proteins [8]. They are typically expressed in a cell-specific, tissue-specific manner [9], and show a relatively low degree of evolutionary conservation [10]. Studies have described the importance of specific features of primary sequence, secondary structure, and genomic position for lncRNA functionality [11]. Another property that expands the functional versatility of lncRNAs is their capacity to associate with DNA, RNA, and proteins [12]. Interest in lncRNAs among members of the scientific community has led to an upsurge in the discovery and characterization of these transcripts in a variety of models and systems, where they have been implicated in diverse biological processes and disease-related pathways [13]. LncRNA involvement has been observed in a variety of sensorineural systems, such as the retina [14, 15] and the olfactory bulb [16, 17], but there have been limited data published about the inner ear [18]. Mouse inner ear RNA-seq data generated by our laboratory revealed the repertoire and abundance of lncRNAs expressed in the sensory organs of the inner ear during development [19]. Considering the widely-known roles of lncRNAs in different systems as well as pathological pathways, identifying and characterizing these molecules and their function becomes essential for inner ear research.

In this study, we present a comprehensive spatiotemporal characterization of four lncRNAs expressed in the inner ear. Our results demonstrate a cell-specific pattern of expression for each lncRNA, established for the first time for inner ear lncRNAs using RNAscope. We further analyzed the sub-cellular localization and expression patterns of these lncRNAs in cells and different mouse tissues, respectively. Furthermore, we suggest a potential mechanism through which *Gas5* may exert its regulatory role in the *Notch1* pathway.

## Materials and methods

### Mouse to human lncRNA LiftOver

Mouse lncRNA coordinates (mm10) were converted to human (hg19) using the UCSC LiftOver tool. The human genomic loci were next intersected with human deafness genomic loci from the Hereditary Hearing Loss Homepage [20] using BEDTools.

### Mouse inner ear dissections

All procedures involving mice met the guidelines described in the National Institutes of Health Guide for the Care and Use of Laboratory Animals and were approved by the Animal Care and Use Committees of Tel Aviv University (01-16-100 and 01-17-033) and the NIH (Protocol number 1235). At Tel Aviv University, C57BL/6J mice were purchased from Envigo, Rehovot, Israel or from the Tel Aviv University Meyers Facility for Transgenic Modeling of Human Disease. At the NIH, mice were obtained from the Jackson Laboratory and housed in the Porter Neuroscience Research Center Shared Animal Facility.

For the isolation of postnatal tissue, newborn mice were euthanized by decapitation. Embryonic tissue was isolated by euthanizing pregnant females with CO_2_, followed by immediate decapitation of the embryos. The head skin was removed and the dorsal part of the skull was opened along the midline. The brain tissue was removed and the auditory nerve was pulled out and cut. The inner ear was removed from the temporal bone, and placed in phosphate buffered saline (PBS, Biological Industries). The cochlea was further dissected, and the sensory epithelium was separated, base to apex, from the spongy modulus bone and immediately placed in RNA*later®* (Sigma-Aldrich).

### RNAscope™

LncRNA candidate probes were designed and purchased from Advanced Cell Diagnostics. CD1 E16.5 and newborn mouse ears were dissected and fixed in 4% paraformaldehyde (PFA) for 24 h at 4°C. The ears were placed in a 5%-30% sucrose gradient (the concentration of sucrose was increased when the tissue sank inside the tube) at 4°C. The ears were left to sink in 30% sucrose again, and were then incubated overnight at 4°C in 30% sucrose with the addition of 2 drops of optimal cutting temperature (OCT) compound. The next day, the ears were transferred to a 1:1 concentration of 30% sucrose and OCT compound and shaken for 1 h, before being added to OCT compound, frozen on dry ice, and stored at −80°C. The ear molds were cryo-sectioned to produce 10 μm sections, which were mounted on slides. Probe hybridization was according to the manufacturer’s protocol and images were collected using a Zeiss LSM 710 microscope (40x and 63x magnifications).

### Cell culture

N2a cells were cultured in Dulbecco’s modified Eagle’s medium (DMEM) supplemented with 10% fetal bovine serum (FBS) and 1% penicillin-streptomycin solution (Biological Industries).

### RNA isolation from mouse cochlea

Total RNA was isolated from mouse cochlea sensory epithelia using the Direct-zol^TM^ RNA MiniPrep (Zymo Research). The RNA was quantified using a Nano-drop^®^ (Thermo Fisher Scientific).

### Subcellular fractionation of cells

To determine the cellular localization of lncRNAs, cytosolic and nuclear RNA fractions were isolated from N2a cells using the PARIS™ Kit (Invitrogen), in accordance with the manufacturer’s instructions. RNA quality was assessed using the Agilent Tapestation system, with a measured RNA Integrity Number equivalent (RIN^e^) score of 10 for both nuclear and cytoplasmic fractions. RNA was also quantified using a Nano-drop^®^ (Thermo Fisher Scientific).

### Quantitative real-time RT-PCR assay

RNA isolated from the cytoplasmic and nuclear fractions of N2a cells, and inner ear tissues, as well as the Mouse total RNA Master panel (Clontech), were used to study the level of expression of candidate lncRNAs. Aliquots of 500 and 750 ng of RNA were used to synthesize single-stranded cDNA from different mouse tissues and the cytoplasmic and nuclear fractions of N2a cells, respectively. Reverse transcription of RNA was performed using the qScript reverse transcription kit (Quantabio). The cDNA was kept at −20°C. Expression levels of protein coding genes and lncRNAs were evaluated with the PerfeCTa SYBR^®^ Green FastMix^®^ ROX kit (Quantabio). *Gapdh* was used as an internal control. Primers were designed using the Primer3 online tool (https://www.bioinfo.ut.ee/primer3-0.4.0/) and ordered from IDT. Primer sequences are provided in Supplementary Table 1. Reactions were carried out in the StepOne^TM^ Real-Time PCR System (Thermo Fisher Scientific) and the analysis was performed using StepOne^TM^ software v2.1. At least three independent experiments were conducted with duplicates and a no-template control (NTC) as a negative control. Melt-curve analysis was done in every reaction for the confirmation of a single product. Data was analyzed by the relative quantification (ΔΔCt) method and expressed as mean ± standard error of the mean (SEM). For the fractionation analysis, the results were calculated as initial quantity, presented as cytoplasmic vs. nuclear fractions, mean initial quantity ±SEM.

### Analysis of single cell data

Expression of each lncRNA was examined in the data sets generated by [21] using Seurate v3. Existing Seurat objects were queried for expression of each lncRNA and the results were presented using the Dotplot function.

## Results

### Selection of functionally relevant inner ear lncRNAs

With the aim of identifying functionally relevant inner ear lncRNAs, we integrated our previously published RNA-seq data with GENCODE (Encyclopedia of genes and gene variants, https://www.gencodegenes.org/) annotations, which includes protein-coding loci with alternatively spliced variants, non-coding loci, and pseudogenes [19]. Approximately 3,700 lncRNAs of different types were found. As the next step, we established a set of criteria for selecting functionally relevant lncRNA candidates for further analysis. These included three obligatory criteria: high expression in the inner ear, conservation in the human transcriptome (see below), and above-threshold expression in at least two well-characterized mouse cell lines. We selected cell lines derived from diverse tissues in order to facilitate an understanding of the functions and modes of action of the lncRNAs. The cell line options chosen for this purpose were 3T3, a mouse fibroblast cell line; B16, a mouse melanoma cell line; and Neuro2A (N2a), a mouse neuroblastoma cell line. These criteria enabled us to study the lncRNAs in both inner ear tissue and cultured cells, thereby expanding our options for analysis.

Since many lncRNAs act in *cis* to regulate the expression of nearby genes [22], we examined the genomic context of the lncRNAs in order to assess any potential association to deafness. Whether the lncRNAs overlapped syntenic chromosomal locations of human deafness loci was a consideration, but was a non-obligatory criterion. Sixteen lncRNAs were selected according to our criteria (Supplementary Table 2). A UCSC track illustrating the overlap between *Gas5* and a deafness locus, DFNA7, is displayed (Supplementary Figure 1). This list was further processed in order to eliminate lncRNAs with repetitive sequences or lncRNAs that host several small nucleolar RNAs (snoRNAs) or microRNAs (miRNAs), as these are more challenging to study. The lncRNAs *Gas5*, *1810014B01Rik,* and *2700046G09Rik* were chosen for further analysis (Table 1). An additional lncRNA was selected because of its unique chromosomal location and previously reported association to DNA methylation dynamics in the inner ear [23]. *Xloc_012867* is located in the region near *Gjb2*, a prominent deafness-related gene [24, 25] (Table 1). The different isoforms of *Xloc_012867* expressed in the inner ear are presented in Supplementary Figure 2, as a multiple sequence alignment.

**Figure 1.**
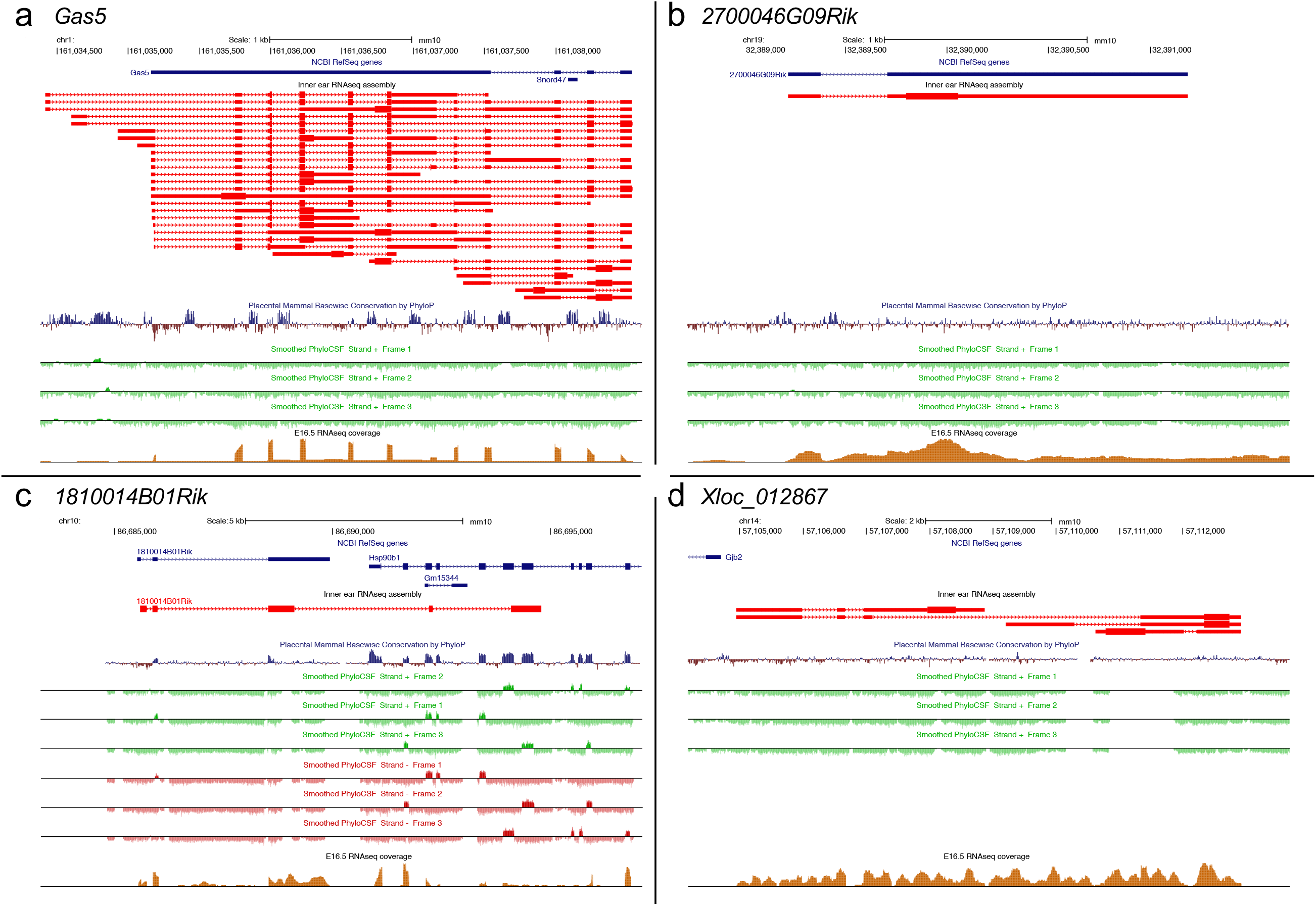
Genomic characterization of lncRNA candidates. UCSC Genome Browser tracks depicting mouse gene variants (red) with PhyloP conservation tracks shown below (dark blue- conserved, red- not conserved). PhyloCSF results presented in green (positive strand), show negative coding potential for all three frames available. Bottom panel tracks indicate mouse inner ear RNA-seq coverage. An E16.5 cochlea sample is presented as an example. The rest of the samples demonstrated the same pattern of expression. The thick boxes in the PLAR tracks represent the longest ORFs of each variant. (A) *Gas5*, (B) *2700046G09Rik,* (C) *1810014B01Rik* (also presenting the negative (red) strand for PhyloCSF), and (D) *Xloc_012867.*

In order to identify lncRNAs involved in regulating important pathways in hearing development and maintenance, we employed a number of approaches to characterize the spatiotemporal expression patterns of the lncRNA candidates, as described below.

**Table 1.** lncRNA gene candidates.

| GENCODE symbol | Mouse cell line expression | Co-located human deafness gene/ locus | Distance from deafness gene/ locus |
| --- | --- | --- | --- |
| *Gas5* | 3T3, B16, N2a | DFNA7* | Overlaps |
| *1810014B01Rik* | 3T3, B16 | *SLC17A8* | 3.5 Mb |
| *2700046G09Rik* | 3T3, B16, N2a | *PCDH15* | 3.1 Mb |
| *Xloc_012867* | None | *GJB2* | 4.4 Kb |

\*A known deafness-associated locus with no identified gene (DFNA: DeaFNess with an autosomal dominant mode of inheritance).

### Computational analysis of lncRNAs

Computational tools were employed to conduct a thorough characterization of genomic loci, conservation and coding potential of each of the lncRNA candidates. The different lncRNA variants expressed in the inner ear are found in the output of the pipeline for lncRNA annotation from RNA-seq data (PLAR) tracks (Fig. 1). The PLAR method was employed for the identification of lncRNAs as part of the computational process presented in a paper published by our group [19]. Since vertebrate lncRNAs are on average approximately 1000 nt long, multiple putative open reading frames (ORFs) can be expected to occur by chance within their sequences [26, 27], and we visualized the longest ORF in each sequence.

*Gas5* (growth arrest specific 5) is a well-known long intergenic non-coding RNA (lincRNA) located on chr1:161,034,422- 161,038,539 (mm10 assembly). It has 25 known variants in the inner ear, with lengths ranging between 759 and 4,118 bp. *Gas5* exhibits no significant evolutionary sequence conservation among mammals, as measured by PhyloP, an algorithm that analyzes and scores the evolutionary conservation at individual bases [28]. Conserved sequences located inside *Gas5* introns mark the location of snoRNAs, which are conserved between mice and other mammals, as well as other vertebrates (Fig. 1A). *Gas5* was first identified due to its preferential expression in the growth arrest phase of the cell cycle. Lacking the properties of a coding gene, it was first characterized as a snoRNA host, but has now been defined as a lncRNA [29] with functions including the control of apoptosis [30], and the maintenance of mouse embryonic stem cell (mESC) self-renewal and proliferation by inhibiting endodermal lineage differentiation [31]. As functional micropeptides may lay hidden within RNAs that are annotated as putative lncRNAs [32], we examined the coding potential of *Gas5* using PhyloCSF, which can discriminate between protein coding and non-coding transcripts based on their evolutionary signature [33]. This analysis confirmed that *Gas5* has no coding potential, as the PhyloCSF score was negative in all three possible reading frames (Fig. 1A).

*2700046G09Rik* is an annotated lncRNA located at chr19:32,389,216-32,391,184 (mm10 assembly) with no significant sequence conservation among mammals and no recognizable coding potential, evaluated by PhyloP and PhyloCSF, respectively (Fig. 1B). It has a single isoform expressed in the inner ear, with a length of 1,969 bp. *2700046G09Rik* was shown to be involved in a mechanism responsible for regulating myelin production in oligodendrocytes [34]. miR-23a was presented as a negative regulator of *Pten* and *LamnB1* and a positive regulator of the expression of *2700046G09Rik* [35].

*1810014B01Rik* is a lncRNA located at chr10:86690209- 86701968 (mm10 assembly). It has a single isoform expressed in the inner ear, with a length of 9,206 bp. *1810014B01Rik* shows a certain amount of conservation along the gene sequence. PhyloCSF scores were negative in all three frames, except for some short sequences (Fig. 1C), which correspond to regions potentially coding a recent characterized BRAWNIN peptide (see below). Moreover, some regions with positive scores match the overlapping coding gene and the antisense strand positive PhyloCSF scores. *1810014B01Rik* was proposed as a potential late biomarker of Alzheimer’s disease in a transgenic mouse model for the disease [36].

*Xloc_012867* is a novel lncRNA located at chr14:57104950-57119212 (mm10 assembly). There are four different isoforms expressed in the inner ear, with lengths varying between 2,293 and 7,963 bp. *Xloc_012867* is located in the intergenic region between *Gjb2* and *Gjb6*. There is no significant conservation among mammals and the PhyloCSF scores were negative in all three frames (Fig. 1D). *Xloc_012867* was first identified in our RNA-seq data of mouse inner ear [19] and was also described in our study on the mouse inner ear methylome [23].

The overall rapid sequence turnout displayed by all four lncRNAs, despite the fact that all four are conserved in human (see below), is characteristic of their class members. This, together with their lack of substantial coding potential, and other unique characteristics, made them strong candidates for further analysis in the inner ear.

### Inner ear lncRNA expression analysis

With the aim of learning more about the candidate lncRNAs expression in the inner ear we used RNAscope^®^ (Advanced Cell Diagnostics, ACD), an *in situ* hybridization approach, which has the advantages of higher specificity and higher signal to noise ratio [37]. This technique is based on a unique design of a set of 10-20 Z-shaped pairs of probes, which bind adjacent sequences on the RNA of interest, forming a platform-like structure. A preamplifier, which can only bind when two Z probes are bound, serves as a scaffold for amplifiers to bind, which in turn become scaffolds for the fluorescent probes to bind. Each set of probes provides binding sites for hundreds of fluorescent probes, dramatically amplifying the fluorescence on a single molecule. We assayed the expression of *Gas5*, *Xloc_012867* and *2700046G09Rik* in the sensory epithelia of CD1 mice at embryonic day 16.5 (E16.5) and postnatal day 0 (P0). Figure 2 presents the results in the basal turn, which is the most informative of the three cochlear turns. The mosaic pattern of the hair and supporting cells, formed during inner ear differentiation, begins near the base of the cochlea and continues toward the apex [38]. As the basal duct differentiates at an earlier age than the apex, presenting images of the basal duct enables a broader view of gene expression at each of the ages examined. The results validated the expression of our candidate lncRNAs in the inner ear at both E16.5 and P0. E16.5 samples displayed a similar pattern for all three lncRNAs, with a higher level of expression in the floor of the duct, including both the developing organ of Corti and the adjacent Kolliker’s organ and lesser epithelial ridge cell populations, and a modest level of expression in all other regions of the duct. For both *2700046G09Rik* and *Xloc_012867*, we observed a more diffused expression pattern at P0 compared to E16.5, although the tendency to higher expression in the greater and lesser epithelial ridge cell populations was preserved. *Gas5* was expressed at relatively higher levels at both time points, but with the same patterns of expression observed for the other two lncRNAs (Fig. 2). To confirm and expand our characterization of lncRNA expression, we examined expression at E14, E16, P1 and P7 in a recently published cochlear single cell RNA-seq dataset [21]. Dot plots indicate strong expression of *Gas5* and more limited expression of 2*700046G09Rik* and *1810014B01Rik* throughout cochlear development (Fig. 3). Consistent with the RNAscope results, expression of *Gas5* at E16 and P1 was generally lower in hair cells and in cells located at a distance from the organ of Corti, such as OC90+ cells.

**Figure 2.**
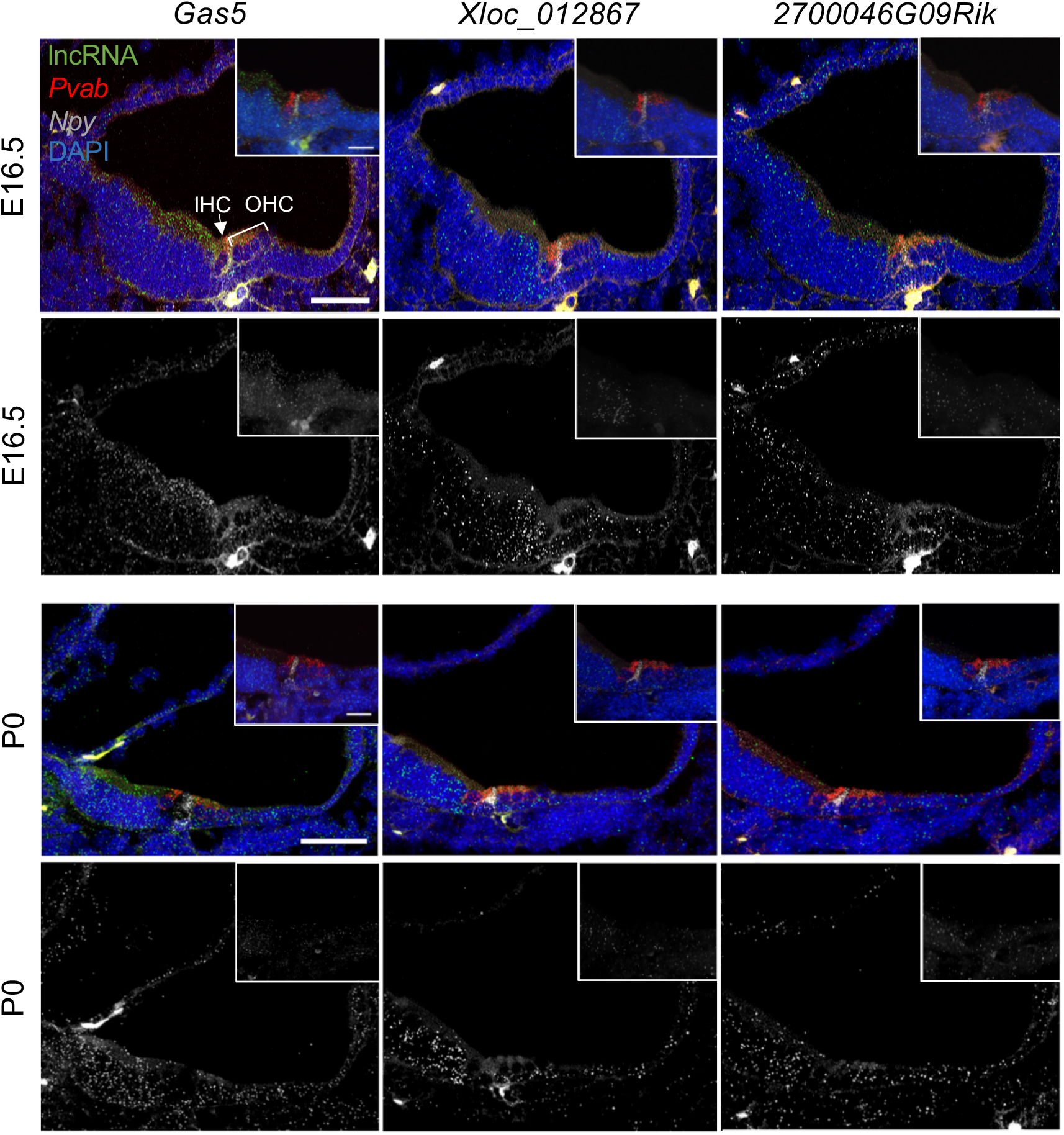
Expression of lncRNAs in the inner ear. E16.5 (upper panel) and P0 (bottom panel) mouse inner ear sensory epithelia samples examined for *Gas5*, *Xloc_012867* and *2700046G09Rik* expression (green). *Pvab* expression (hair cells marker, red), *Npy* expression (inner pillar cells marker, gray) and DAPI (blue) were used as controls. Upper panels for each age indicate all four labels, while the lower panels indicate only the lncRNA expression in grayscale. Insets indicate higher magnification views of the organ of Corti. IHC (inner hair cells), OHC (outer hair cells). Scale bars: main panels = 50 μm, insets = 20 μm.

**Figure 3.**
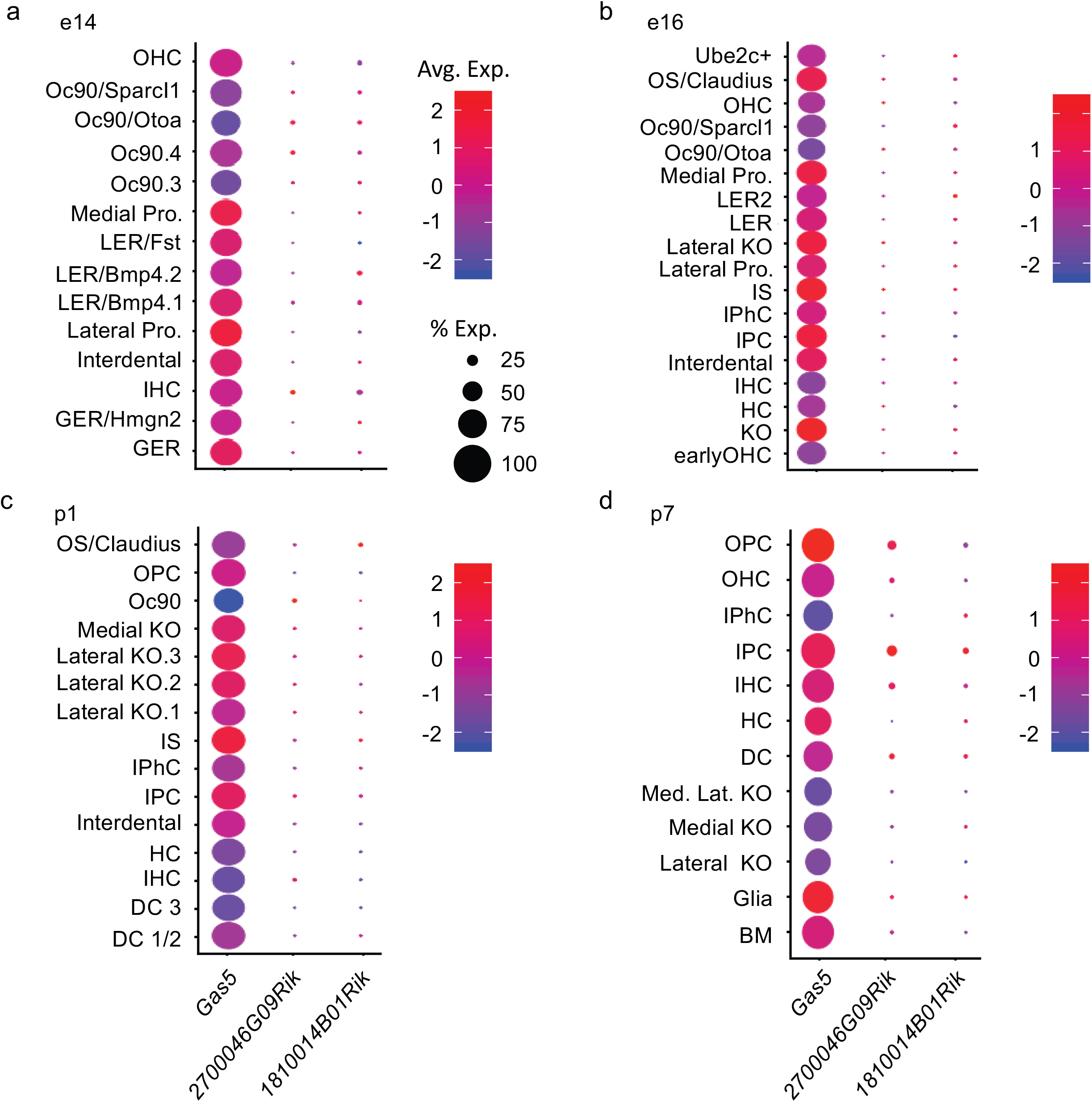
Expression of lncRNA candidates in different cells types of the inner ear. Expression of *Gas5*, *2700046G09Rik*, and *1810014B01Rik* in the indicated cell types based on single cell RNA-seq data [21] at the indicated developmental time points: (A) E14, (B) E16, (C) P1, and (D) P7. Size of each circle indicates percent of cells for each cell type expressing the indicated lncRNA. Color code indicates scaled (z-score) average expression in each cell. BM (basement membrane), DC (Deiters’ cells), DC1/2 (Deiters’ cells rows 1 and 2), DC 3 (Deiters’ cells row 3), GER (greater epithelial ridge), GER/Hmgn2 (greater epithelial ridge / Hmgn2 expressing cells), HC (hair cells), IHC (inner hair calls), IPC (inner pillar cells), IPhC (inner phalangeal cells), IS (inner sulcus cells), KO (Kölliker’s organ cells), Lateral KO (lateral Kölliker’s organ cells), Lateral KO.1 (lateral Kölliker’s organ cells type 1), Lateral KO.2 (lateral Kölliker’s organ cells type 2), Lateral KO.3 (lateral Kölliker’s organ cells type 3), Lateral pro. (Lateral prosensory), LER (lesser epithelial ridge), LER2 (lesser epithelial ridge type 2), LER/ Bmp4.1 (lesser epithelial ridge/ BMP4 type 1 expressing cells), LER/ Bmp4.2 (lesser epithelial ridge/ BMP4 type 2 expressing cells), LER/Fst (lesser epithelial ridge/ Fst expressing cells), Medial KO (medial Kölliker’s organ cells), Med. Lat. KO (medial lateral Kölliker’s organ cells), Medial pro. (Medial prosensory), Oc90 (Otoconin90 expressing cells), Oc90.3 (Otoconin90 type 3 expressing cells), Oc90.4 (Otoconin90 type 4 expressing cells), Oc90/Ota (Otoconin90/ Otoa expressing cells), Oc90/Sparcl1 (Otoconin90/ Sparcl1 expressing cells), OHC (Outer hair cells), OPC (outer pillar cells), Os/Claudius (outer sulcus cells/ Claudius), Ube2c+ (Ube2c expressing cells).

### lncRNA sub-cellular localization patterns

LncRNA sub-cellular localization can range between the nucleus, the cytoplasm, and chromatin [9]. The localization to a specific cellular location may aid in revealing function, as lncRNAs must localize to their place of action. In order to reveal lncRNA localization, we extracted RNA from the nuclear and cytosolic fractions of N2a cells, neuroblastoma cells, which are an accessible source of neuronal tissue and may share common characteristics with inner ear sensory tissue. All four lncRNAs could be detected in both the cytoplasm and nucleus fractions. *2700046G09Rik*, *1810014B01Rik,* and *Xloc_012867* were shown to localize predominantly to the nuclear fraction, while *Gas5* was expressed at relatively higher levels in the cytoplasmic fraction (Fig. 4).

**Figure 4.**
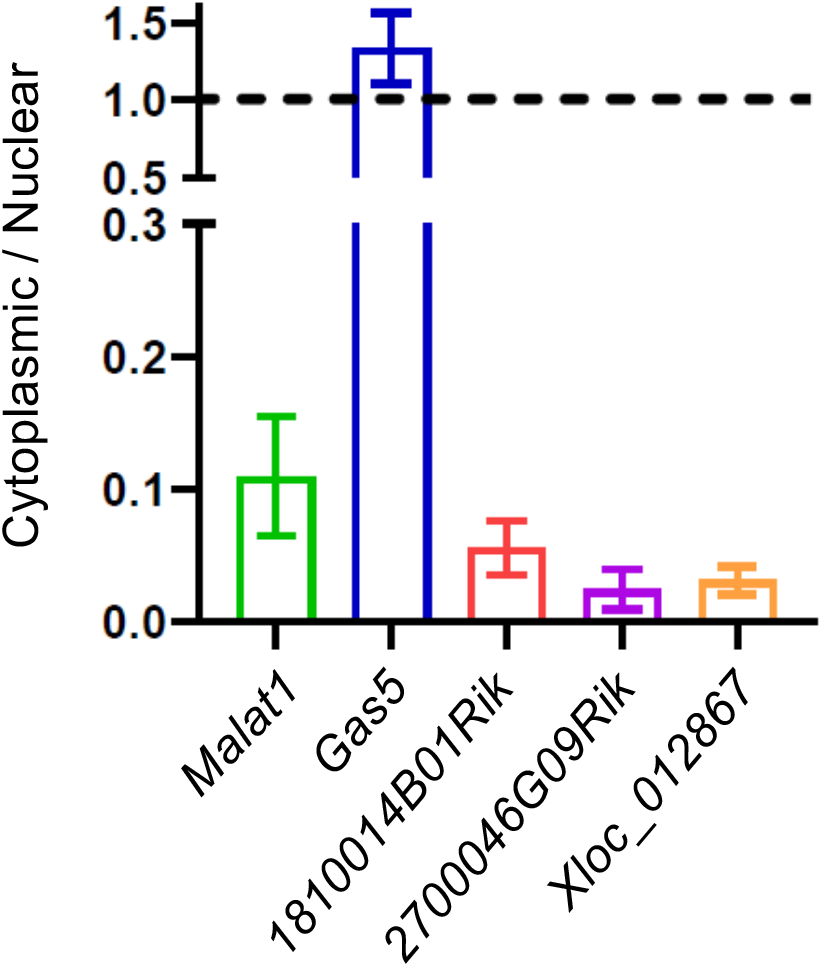
lncRNA sub-cellular localization. IncRNA expression in the cytoplasmic and nuclear fractions of N2a cells measured by qRT-PCR, presented as cytoplasmic vs. nuclear fractions mean initial quantity ±SEM, n=3. *Malat1*, known to be localized to the nuclear fraction, served as a control.

### LncRNA tissue specificity

Exploring the expression patterns of lncRNAs in different tissues can provide information about the function of a specific lncRNA. We examined the expression of our lncRNA candidates in samples that were extracted from four embryonic developmental stages and 11 different mouse tissues, including the sensory epithelium of the inner ear (Fig. 5). The expression of *Gas5* was exceptionally high in the eye, spleen, testis, and cochlea, with a more moderate expression measured in the salivary glands, kidney, and liver. The expression of *1810014B01Rik* was particularly elevated in the testis tissue, with a more moderate expression in the brain, eye, heart, kidney, liver, salivary glands, spleen, and cochlea. Embryonic expression was lower. *2700046G09Rik* was expressed at extremely high levels in the testis, compared to the other tissues examined. This trend is also apparent from the published ENCODE transcriptomic data [39]. The expression of *Xloc_012867* in the cochlea was considerably higher than in the kidney, liver, salivary glands, spleen, stomach, and testis. Other tissues showed even lower expression levels (Fig. 5). These results demonstrate the large variability in lncRNA expression with respect to both the localization and levels.

**Figure 5.**
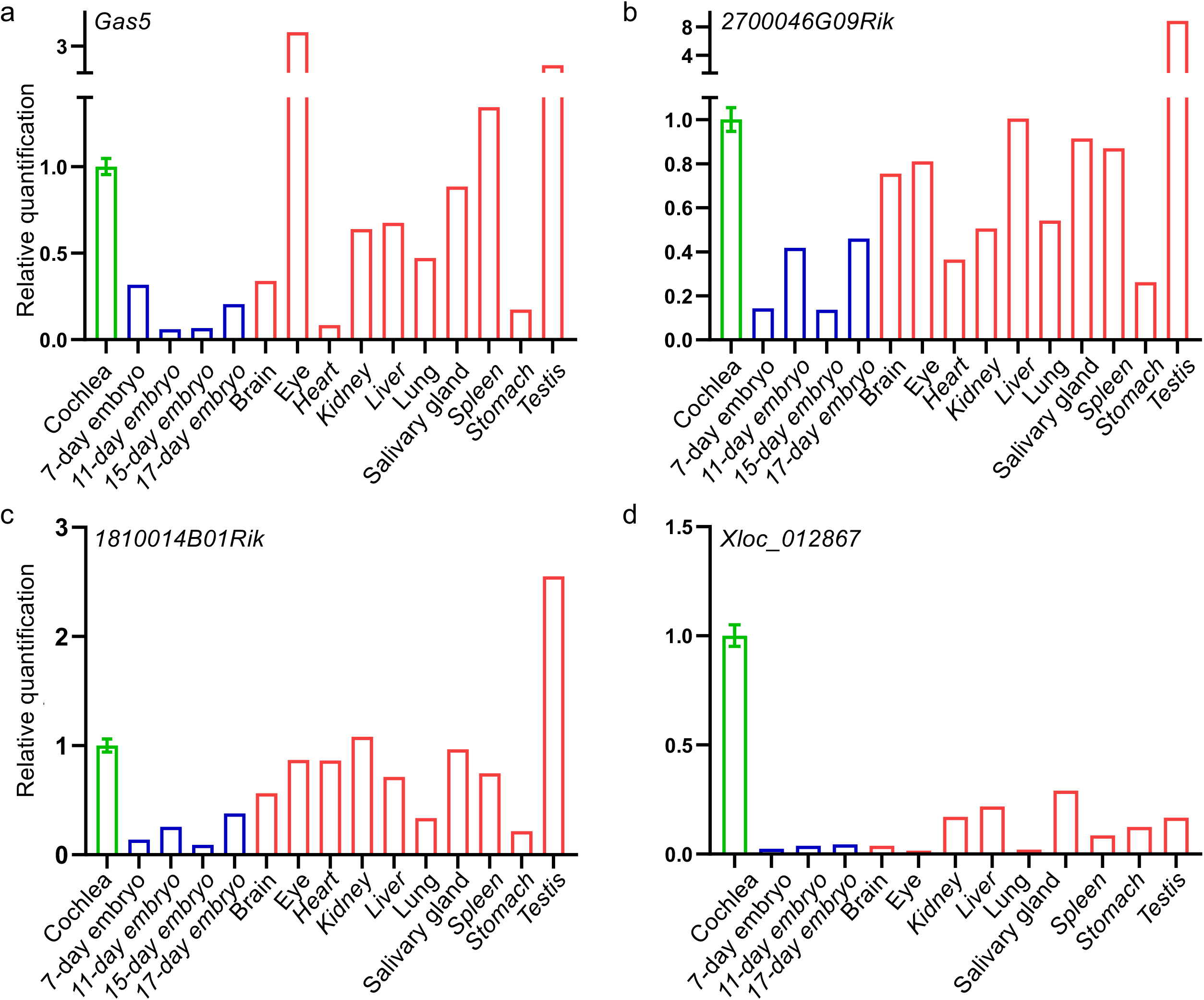
lncRNA expression in a mouse tissue panel, using qRT-PCR. (A) *Gas5*, (B) *2700046G09Rik,* (C) *1810014B01Rik*, and (D) *Xloc_012867.* Mean ± SEM, n=3. Error bars are presented for the cochlea only as the source of the rest of the samples is the Mouse Total RNA Master Panel (Clontech).

## Discussion

Non-coding RNAs (ncRNAs) are crucial for a wide variety of biological processes and deregulation of their expression might lead to various pathologies, including cancer [40], celiac disease [41], and cardiovascular diseases [42], among others. In recent years, advances in technologies have revolutionized our capacity to decode the diverse features of the genome and have enabled the discovery and characterization of different classes of ncRNA molecules, including lncRNAs. These have proved to be a diverse group of molecules with the ability to exert their functions in *cis*, in *trans* [43], in the nucleus [44], or cytoplasm, and even to be able to shuttle between the two compartments [9]. LncRNAs have tissue-specific expression patterns and are dynamically expressed during different developmental pathways in biological processes and pathologies [45, 46]. However, the role of lncRNAs in the auditory system and their contribution to hearing loss-related pathologies remains largely unclarified. Two lncRNAs, *Meg3* [47] and *Rubie* [48] are expressed in the cochlea and vestibule, respectively. A transcriptomic study of the human inner ear, derived from three individuals whose hearing could not be preserved upon tumor removal, described over 7,000 lncRNAs, with 253 differentially expressed between the cochlea and the vestibule [18]. Our group published a comprehensive analysis of the lncRNA landscape in the auditory and vestibular systems during development [19]. In this work, we presented a comparison between the mouse lncRNA catalogue and those annotated in human. Orthology was examined by both sequence similarity and synteny. We identified 139 lncRNAs with sequence similarity and 1,049 syntenic lncRNAs. A subset of 101 lncRNAs shared both sequence similarity and syntenic location [18, 19].

Here we report the comprehensive characterization of *Gas5*, *Xloc_012867*, *2700046G09Rik,* and *1810014B01Rik,* four lncRNAs, each with its own unique set of properties. One of the criteria for choosing the lncRNAs for further study was their conservation in human tissues. *Gas5* has an annotated homolog in human (https://www.ncbi.nlm.nih.gov/gene/60674). *Gas5* overlaps DFNA7, a deafness-associated locus, for which *LMX1A* is a candidate gene [49]. Since lncRNAs act in *cis* to regulate the expression of nearby genes [22], *Gas5* may regulate this gene. *2700046G09Rik* has a synthetic human lncRNA known as *SGMS1-AS1*. It seems to have limited sequence overlap with the mouse lncRNA, but is transcribed from the same region. The general function of this lncRNA is yet to be determined in human, but it is interesting to note its genomic proximity to the deafness gene *PCDH15*. The human *Xloc_012867* homolog is *AL138688.1/RP11-264J4.10*. *RP11-264J4.10* was suggested to be involved in the regulation of *Gjb6*, a deafness-associated gene [23]. Most recently, parts of the human homolog for *1810014B01Rik*, *C12orf73*, was annotated as coding for a 71 amino acid long peptide named BRAWNIN [50]. It was demonstrated, through functional prediction, proteomics, metabolomics and metabolic flux modeling, as essential for respiratory chain complex III (CIII). These are also similar to sequences located in another mouse gene referred to as *1190007I07Rik*. This could be explained by a duplication or rearrangement event at some point during rodent evolution. Depletion of BRAWNIN in human cells impairs mitochondrial ATP production. Deletion of murine *1190007I07Rik* in mouse embryonic fibroblasts (MEF) using CRISPR/Cas9 demonstrated the requirement for BRAWNIN in CIII assembly or stability in mice as well. The *1810014B01Rik* transcript expressed in mouse inner ear is different in length and genomic structure compared to *1190007I07Rik* and further investigation aimed at identifying a functional protein encoded by *1810014B01Rik* is required. Its role as a functional lncRNA has yet to be excluded.

All four lncRNAs show little evidence of evolutionary conservation, which is characteristic of their class, and appear to be non-coding. We describe the use of RNAscope for the detection of lncRNAs in the inner ear for the first time. This advanced *in situ* hybridization technique validated the expression of our candidate lncRNAs in the inner ear at both E16.5 and P0 and demonstrated a cell-specific pattern. Based on these results, we designed and performed experiments aimed at demonstrating the spatial and temporal expression patterns of these lncRNAs, and promoting our knowledge regarding their potential functional roles. Since a robust inner ear cell line is not available, N2a neuroblastoma cells were used, as an accessible resource of cells from neuronal origin that may share common characteristics with inner ear sensory tissue.

The localization of a lncRNA within the cell can provide information about its molecular role and mechanism of action. LncRNAs differ from protein-coding mRNAs in that they have to be present in the cellular location in which they act [9]. For example, lncRNAs located at the site of transcription in the nucleus may be involved in recruitment of transcription factors or chromatin modifying complexes. Nuclear lncRNAs can also regulate gene expression in *trans*, through binding to distant genomic loci, or be involved in alternative splicing. LncRNAs exported from the nucleus to the cytoplasm after transcription, are often involved in regulation through miRNA sponging [51], or engage in translation interference [52]. Subcellular analysis of N2a cells revealed that three of the four lncRNAs examined, (*2700046G09Rik*, *1810014B01Rik* and *Xloc_012867*), were predominantly expressed in the nuclear fraction. *Gas5*, which is known to shuttle between the nucleus and cytoplasm [53], and has different patterns of localization in various types of cells [9], was predominantly expressed in the cytoplasm. These results demonstrate the variability and specificity of lncRNA expression patterns.

The expression of the four lncRNAs in various mouse tissues illustrated another well-known characteristic of lncRNAs, namely high tissue specificity, with differential levels of expression. Interestingly, *Xloc_012867*, selected because of its compelling genomic location, was expressed at considerably higher levels in the cochlea than in the other tissues examined, suggesting its importance in the inner ear. Since this lncRNA is located in a region near a distinct deafness-related genes, *GJB2 and GJB6* [24, 54], it has potential for being involved in its regulation through a *cis*-acting mechanism.

*Gas5* was highly expressed in the ear and eye. Another critical ncRNA family, the miRNAs, are also robustly expressed in these sensory systems. For example, the members of miR-183 family, miR-183, miR-96, and miR-182, were shown to be abundantly expressed and to target essential genes in certain eye and inner ear cell types [55-57]. The similar expression patterns and the established correlation between high expressions levels of miRNAs and a crucial function in these specific sensory systems, supports our hypothesis that *Gas5* has a critical function in the inner ear. *Gas5* was previously associated with a *Notch1*-related pathway, where miR-137 was shown to be a negative regulator of *Notch1,* with *Gas5* acting as a competing endogenous RNA (ceRNA) [58]. Notch signaling is critical for the formation of the organ of Corti’s hair and supporting cells’ mosaic pattern. In addition, Notch defines the development of the prosensory domain of the inner ear and plays a role in its proliferation and regeneration [59, 60]. In fact the mosaic pattern formation is achieved through *Notch1* lateral inhibition mechanism of downregulation in cells that will develop as hair cells, and upregulation in cells that will become supporting cells [61]. Our expression studies using RNAscope, demonstrate enriched expression of *Gas5* in the prosensory domain of the cochlear turn, which is the source of both hair and supporting cells of the organ of Corti [61]. Considering the correlation between lncRNA localization and site of action, we suggest that a *Notch1*–miR-137–*Gas5* interplay is also engaged in the inner ear. This hypothesis is supported by our qRT-PCR results demonstrating an increase in *Notch1* and *Gas5* expression during development, between E16.5 and P0, accompanied by a decrease in the expression of miR-137 between these same time points (Supplementary Methods, Supplementary Table 3, Supplementary Figure 3). Thus, we propose a miRNA negative regulation and a lncRNA positive regulation imposed by its miRNA sponging activity.

In summary, our findings demonstrate the temporal and spatial expression of specific lncRNAs, *Gas5*, *Xloc_012867*, *2700046G09Rik,* and *1810014B01Rik*, in different cell types of the auditory system. Examination of other pathways leads us to hypothesize that *Gas5* might function in the inner ear as a ceRNA to the Notch signaling pathway. While examining the function of lncRNAs remains a challenge due to the lack of a robust inner ear cell line, CRISPR-Cas9 genome editing may provide a means to study their role in the whole animal. Our data provide new information about mouse inner ear lncRNAs and reveal important insights into the regulatory landscape of lncRNAs in the inner ear. As they have in other diseases, lncRNAs may provide new targets for therapy in deafness. In addition, the experimental design of this study may serve as a reference for the analysis of lncRNAs expressed in other sensory systems.

## Acknowledgments

This research was supported by the Israel Science Foundation (grant No. 2033/16 and 1763/20) (KBA), the Ernest and Bonnie Beutler Research Program of Excellence in Genomic Medicine (K.B.A.), the Nella and Leon Benoziyo Center for Neurological Diseases at the Weizmann Institute of Science (I.U.), and funds from the Intramural Program at the NIDCD DC000059 (M.W.K.). K.B.A. is an incumbent of the Drs. Sarah and Felix Dumont Chair for Research of Hearing Disorders. This work was performed in partial fulfillment of the requirements for a Ph.D. degree by Tal Koffler-Brill at the Sackler School of Medicine, Tel Aviv University, Israel. T.K.-B. is supported by a Professor Heinrich Neumann Fellowship for Hearing Health and N.M.-G. is supported by an Edmond J. Safra Fellowship (https://safrabio.cs.tau.ac.il/).

## Contributions

T.K.B., I.U. and K.B.A. conceived the project and along with M.W.K., designed experiments, analyzed the data, and wrote the manuscript, with input from all authors. T.K.-B, N.M.-G., R.E. and I.U. performed the bioinformatics and data analysis. T.K.-B., S.T., A.A., M.B.-C., E.R., M.W.K. and L.K. performed and analyzed the laboratory experiments. All authors have read and approved the manuscript for publication.

## Additional File 1: Table S1 and Figure S1

**Table S1.**
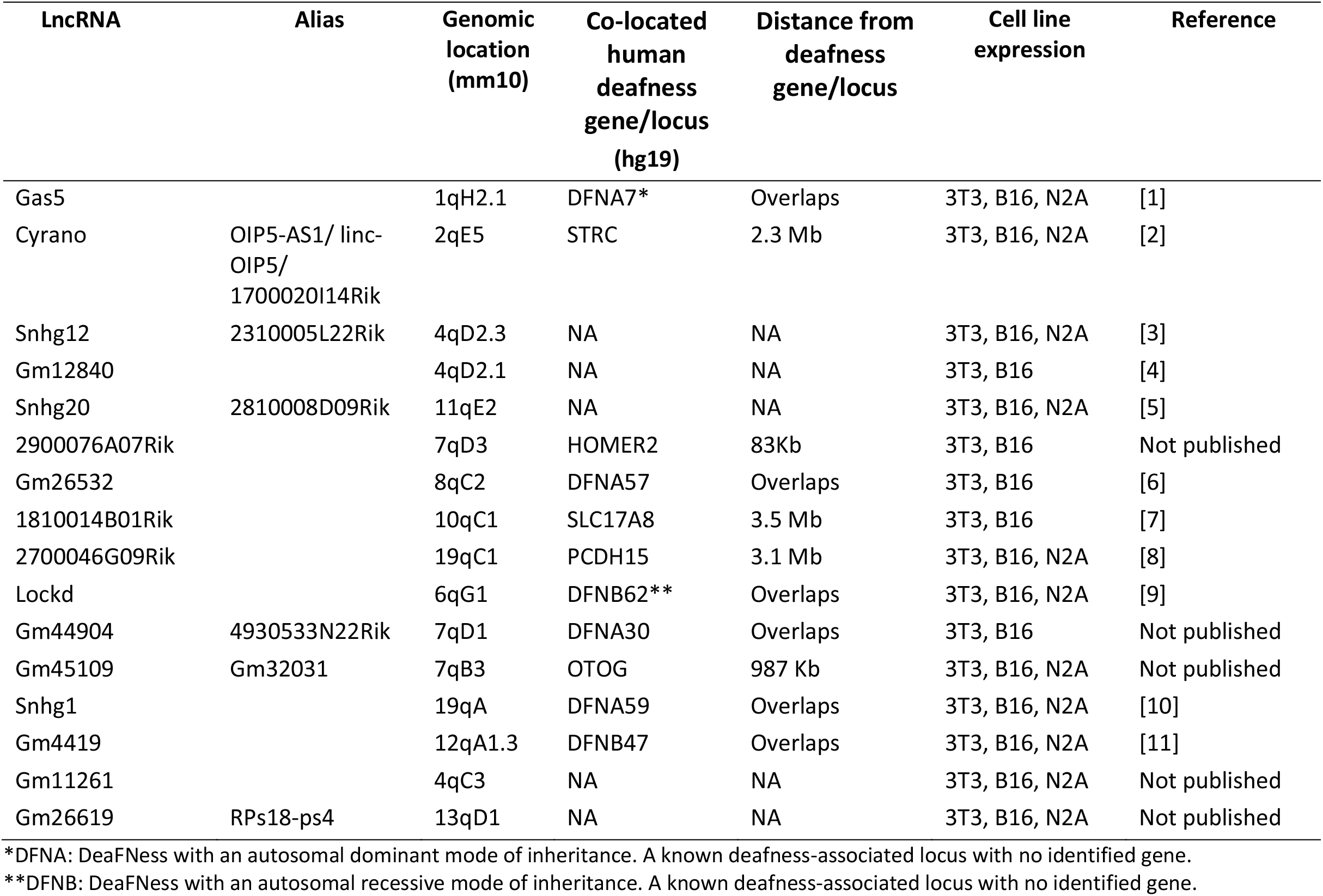
Functionally relevant lncRNA candidates.

**Fig. S1.**
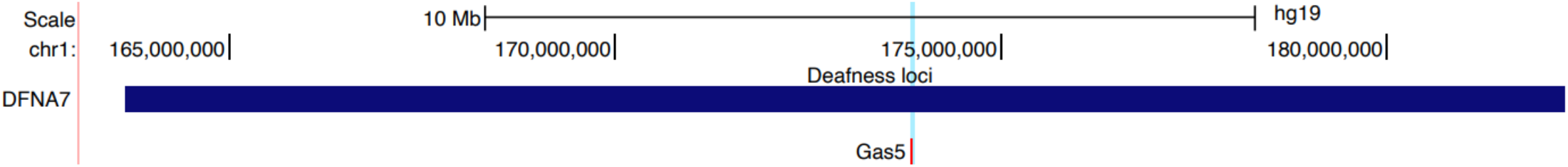
*Gas5* overlap with DFNA7. UCSC Genome Browser track indicating the overlap of *Gas5* with a known deafness locus DFNA7.

---

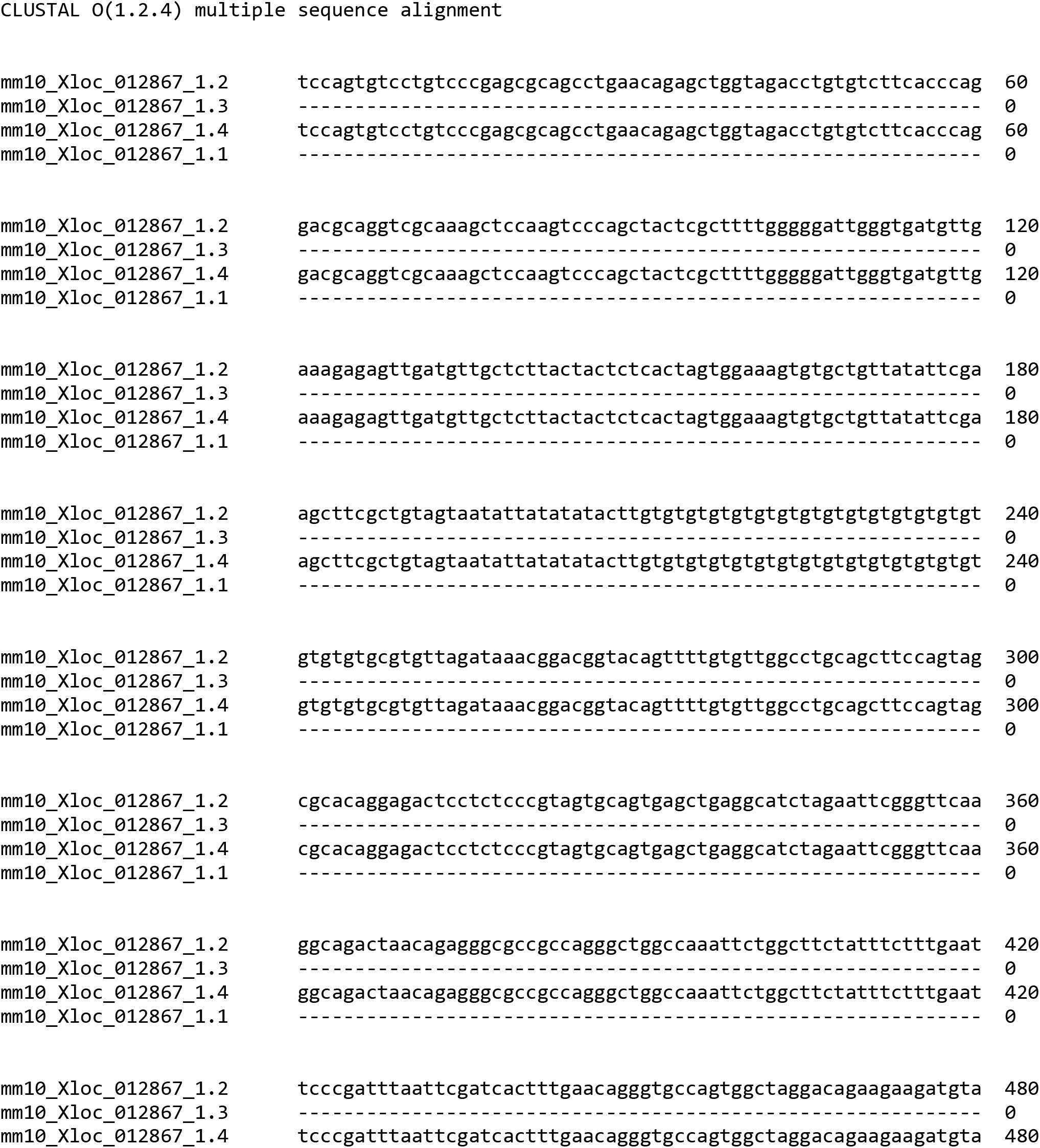
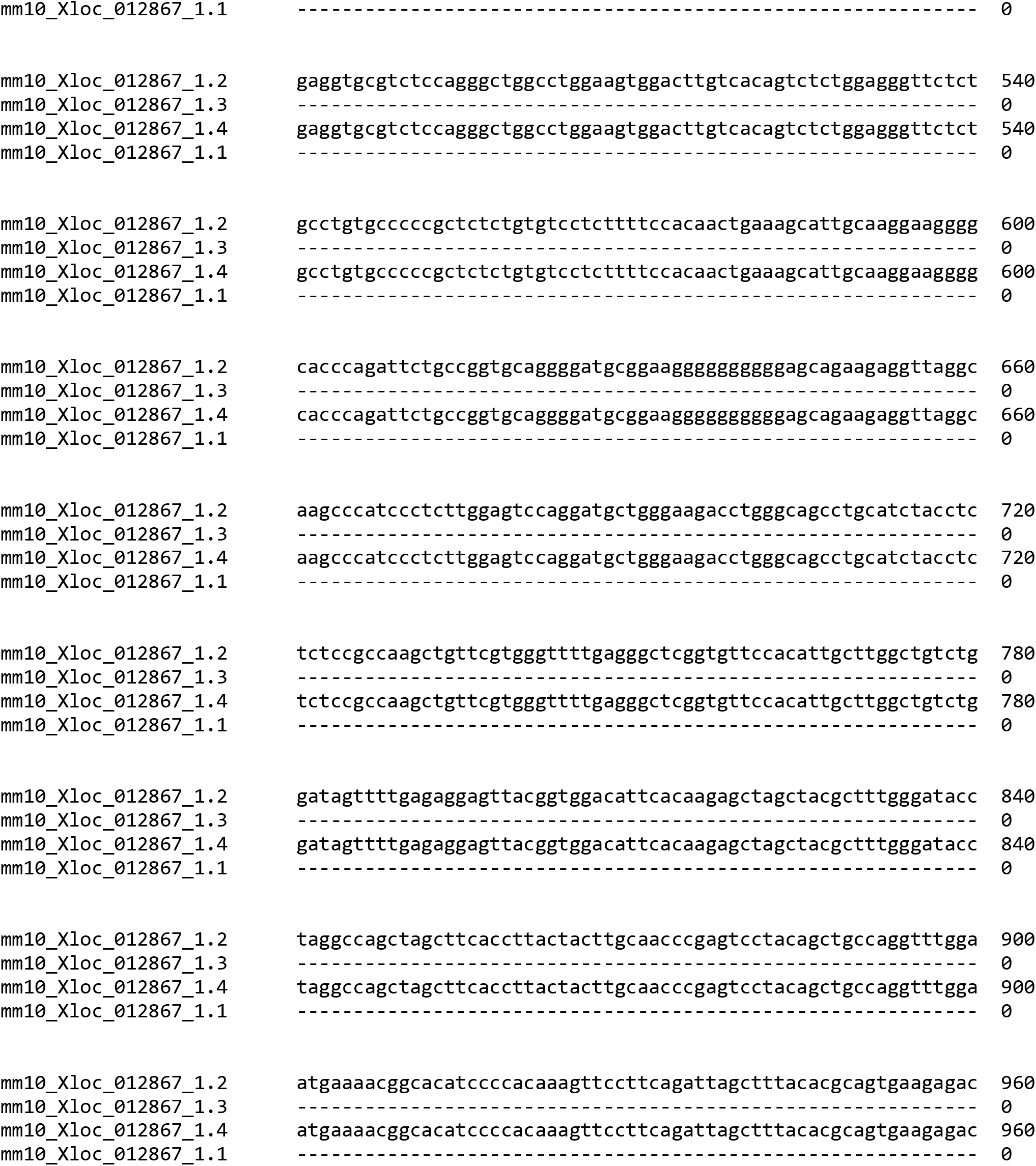
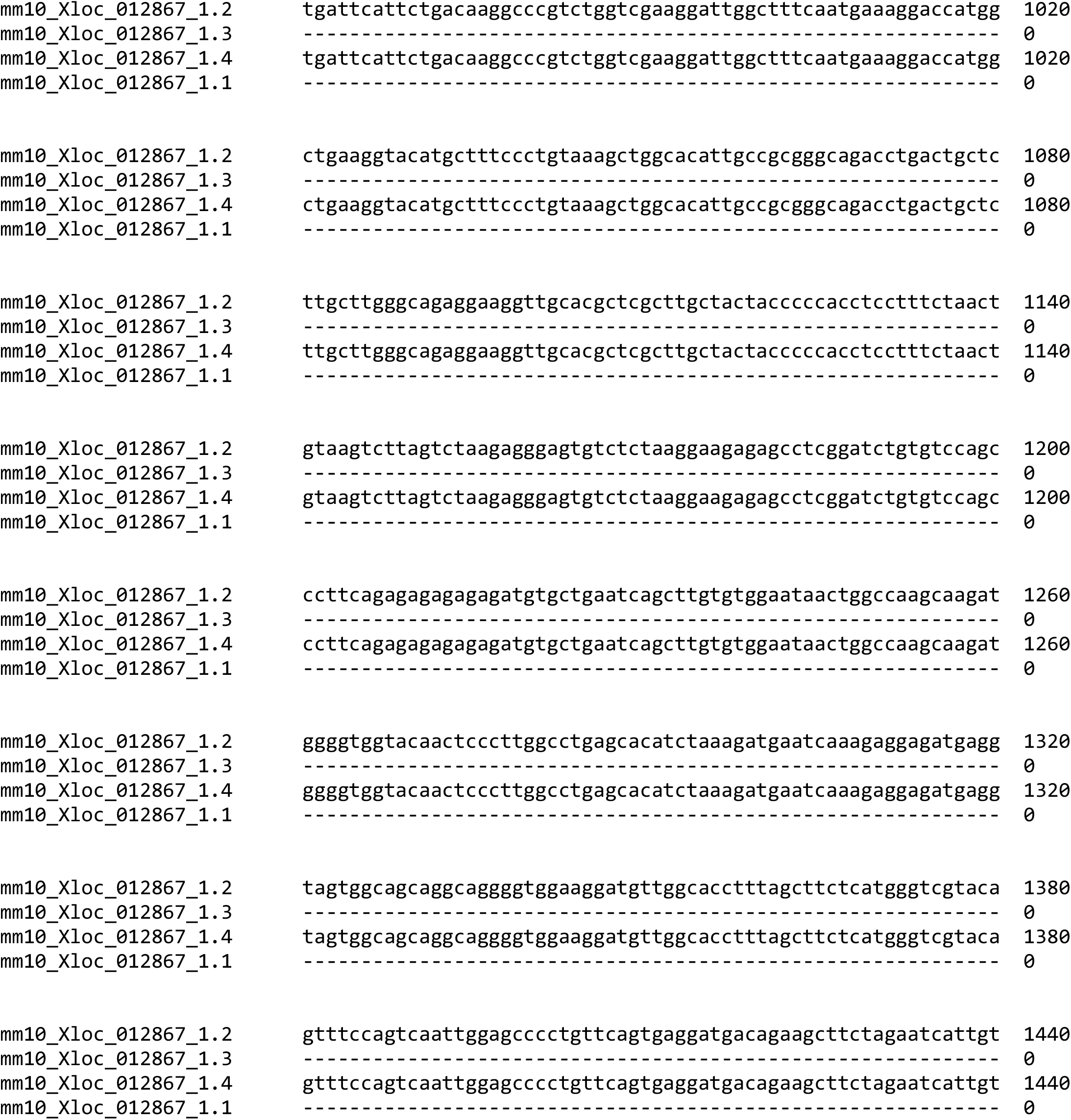
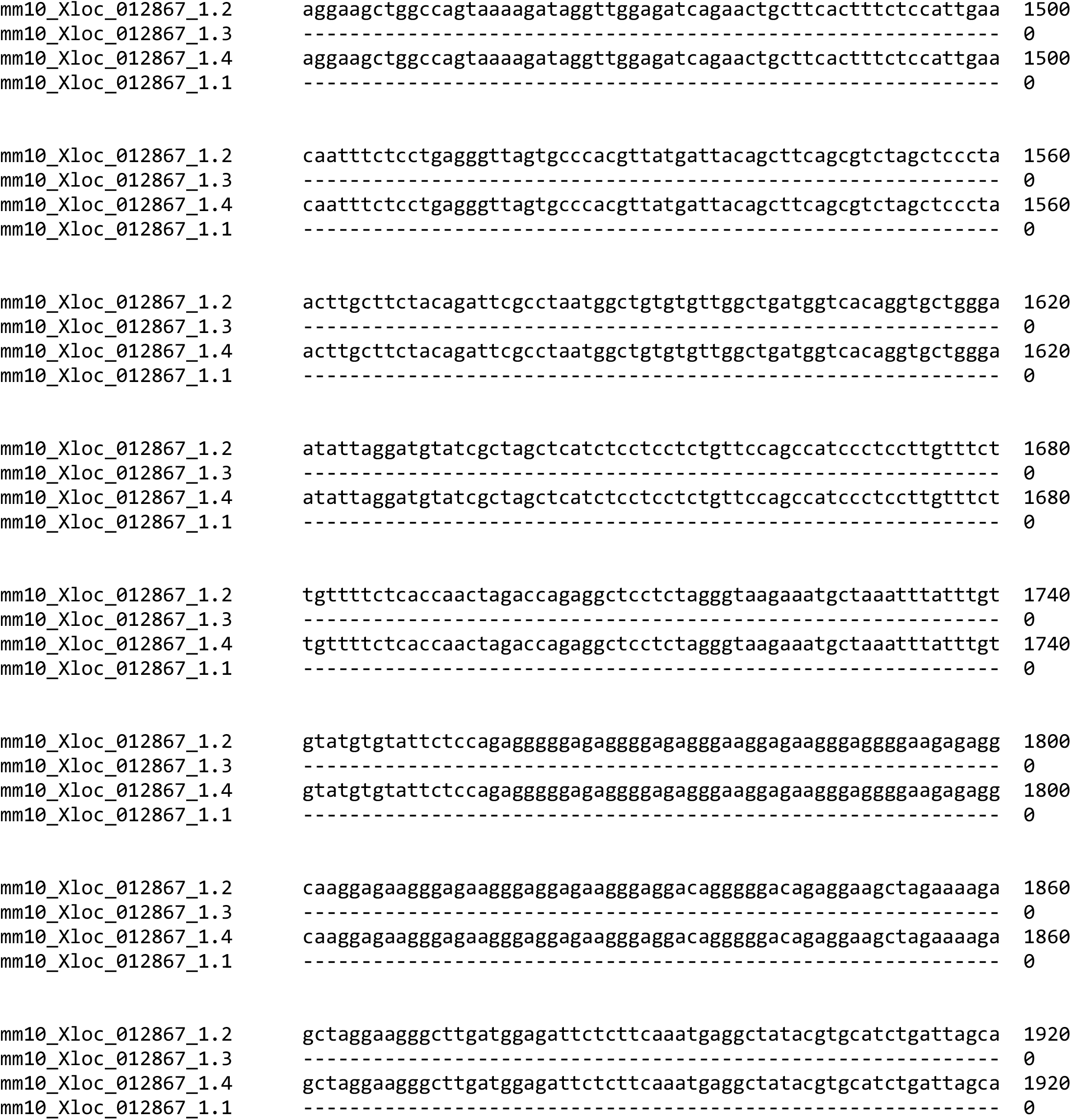
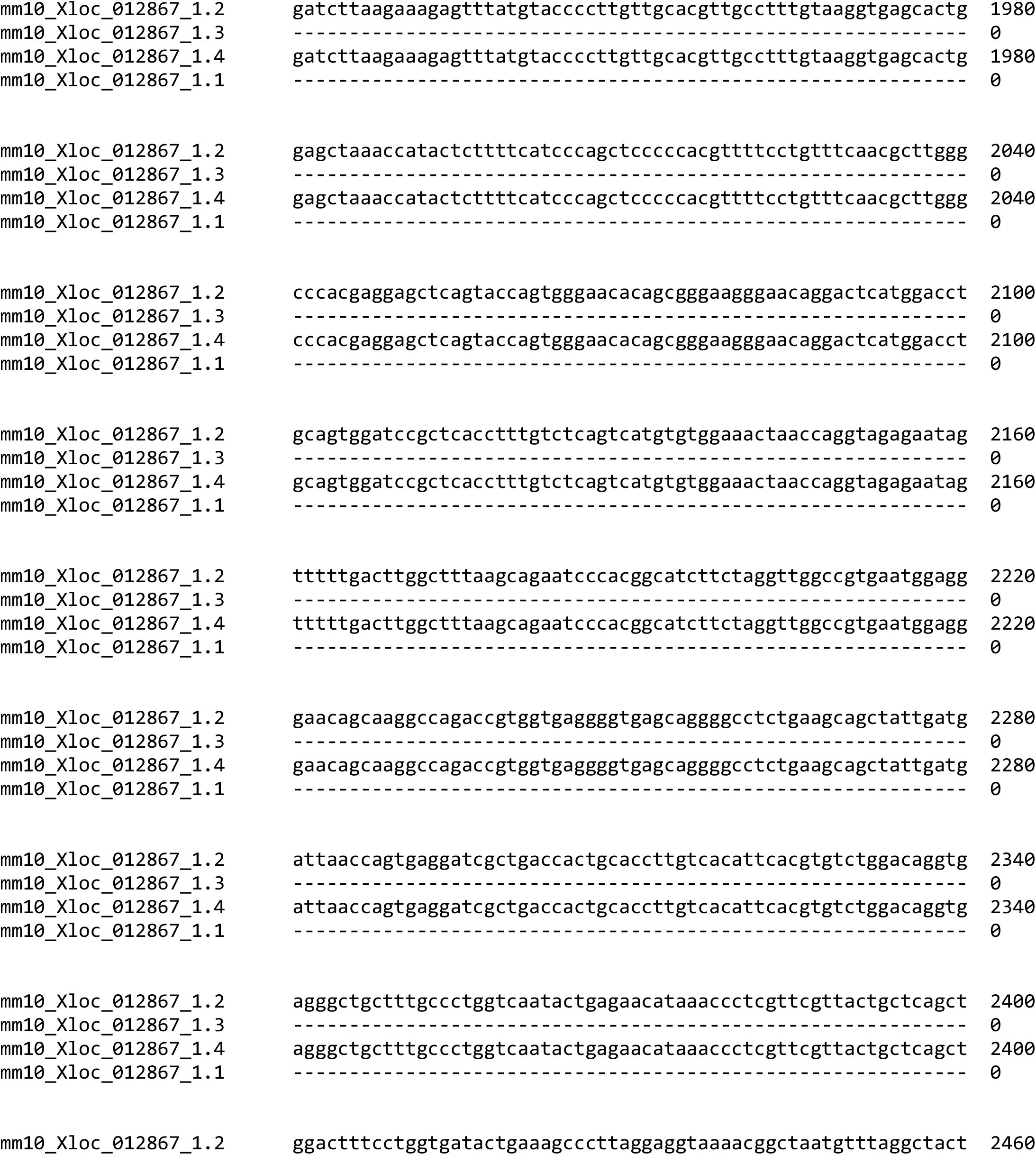
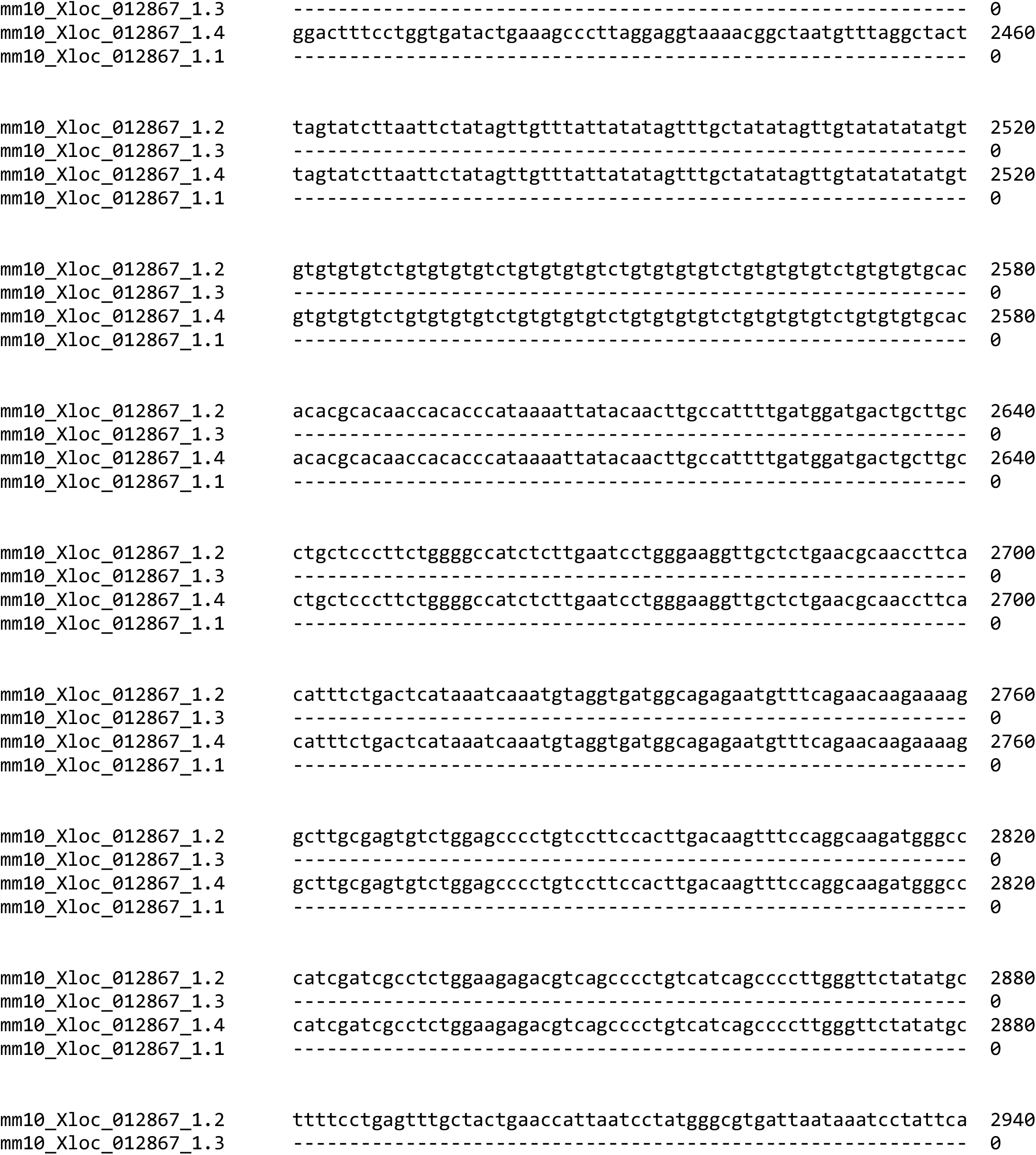
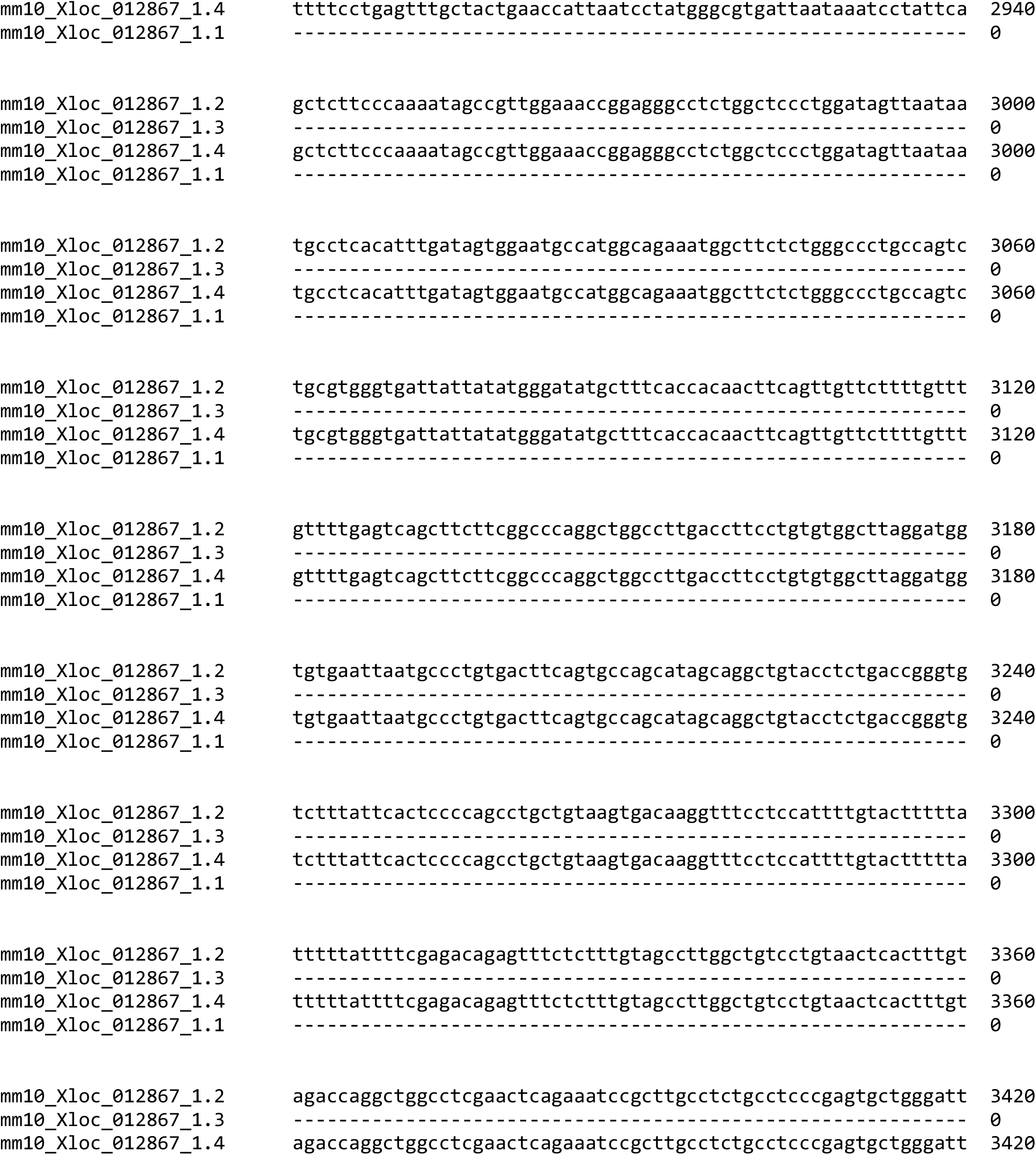
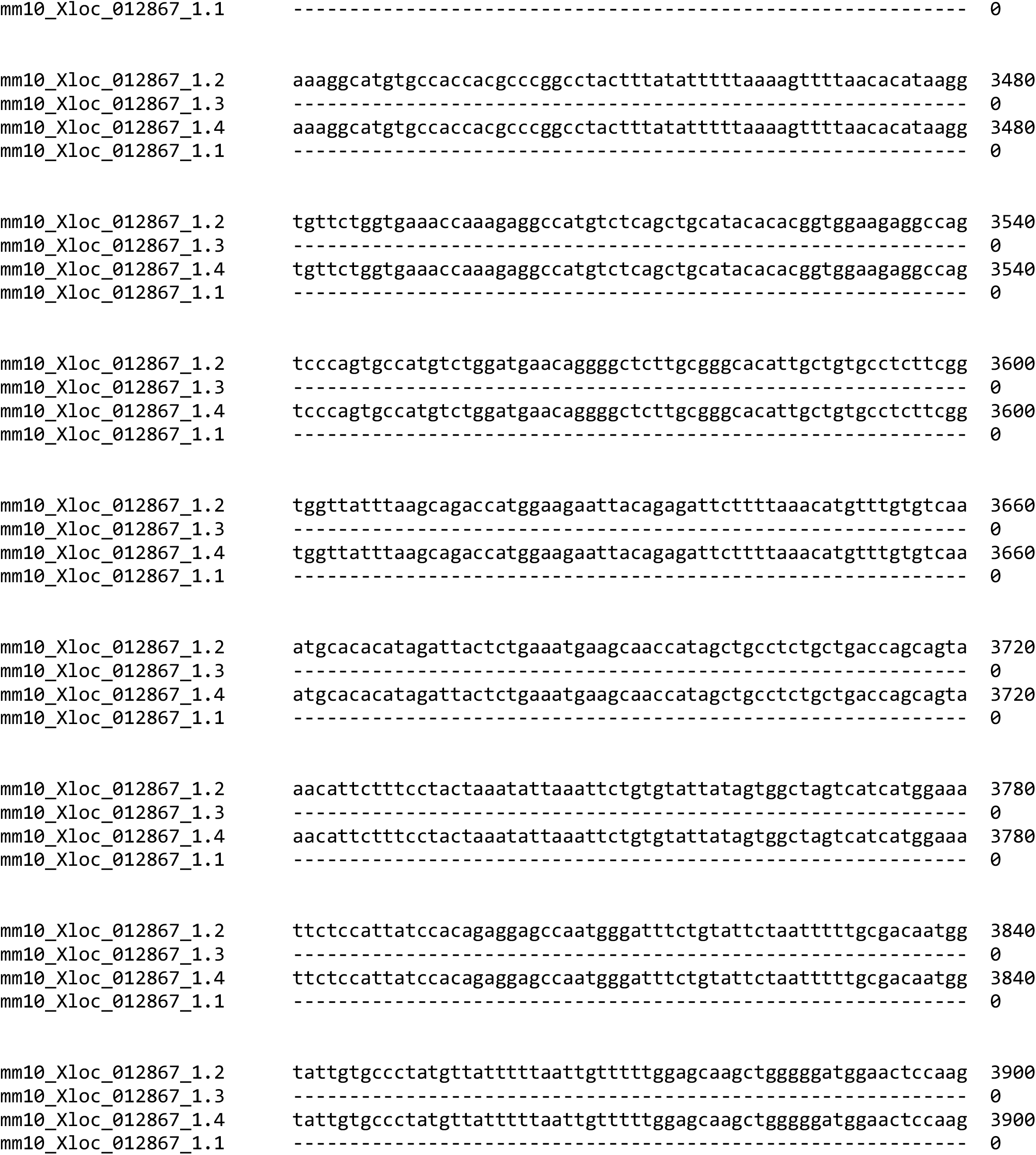
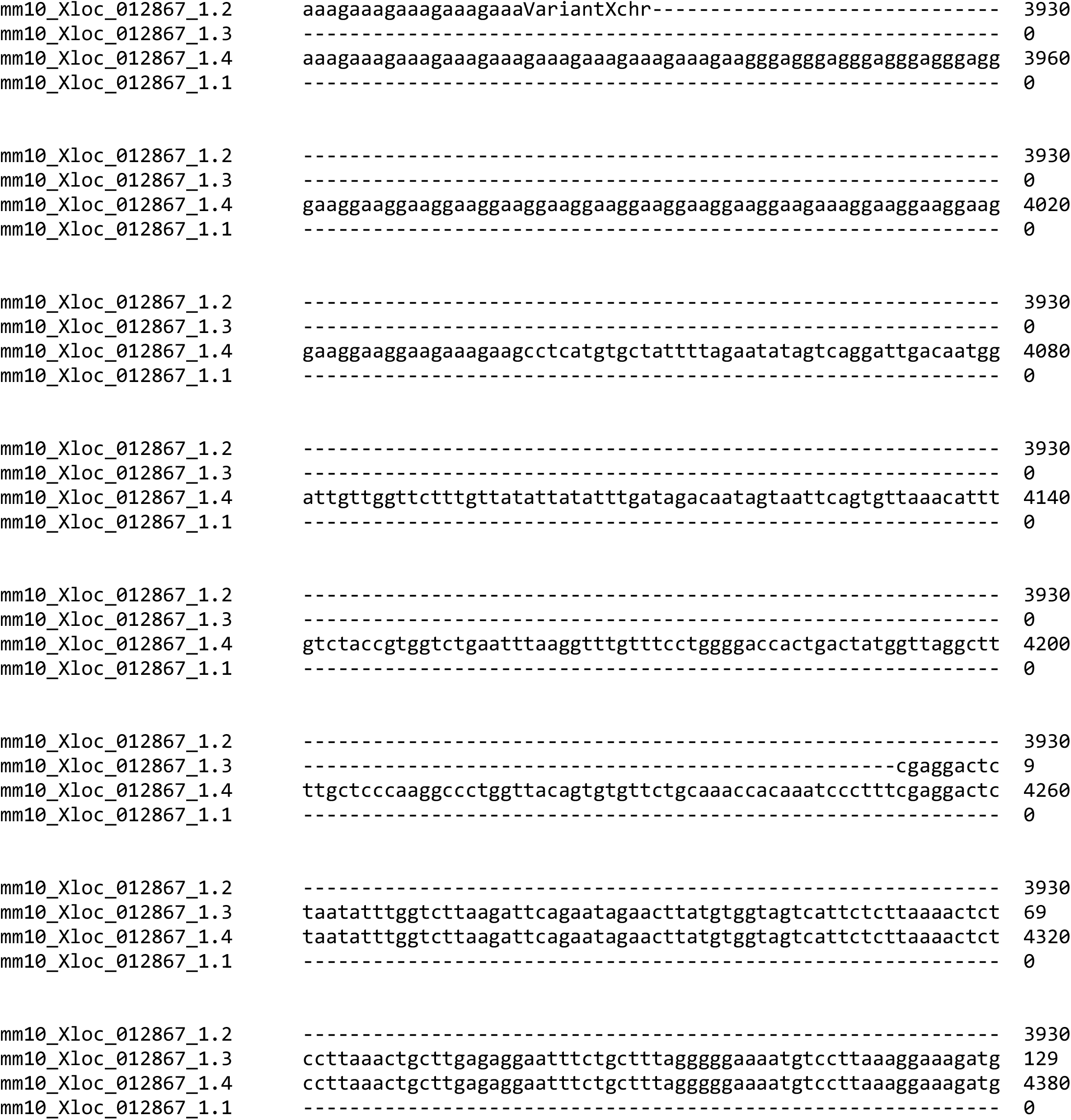
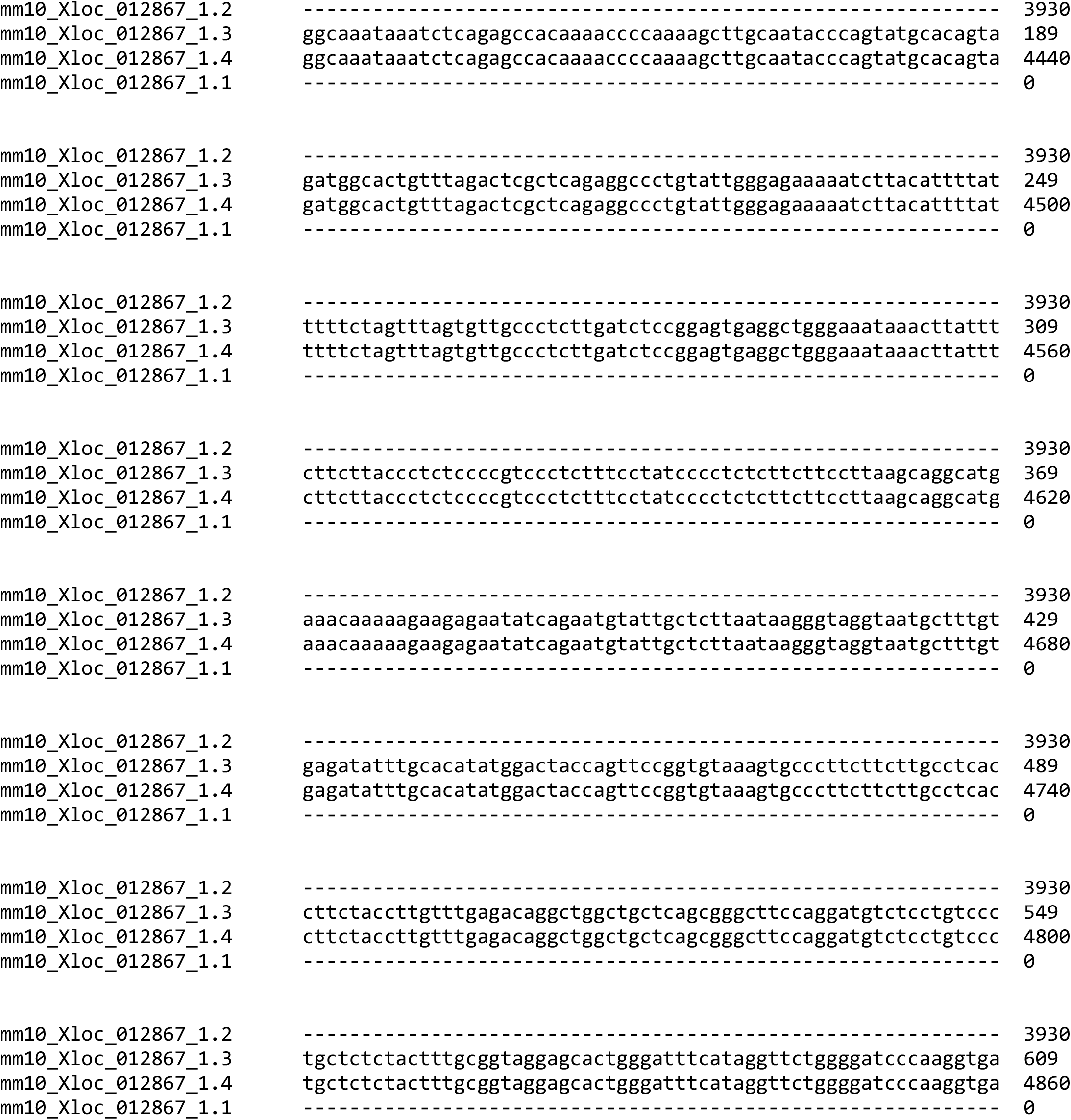
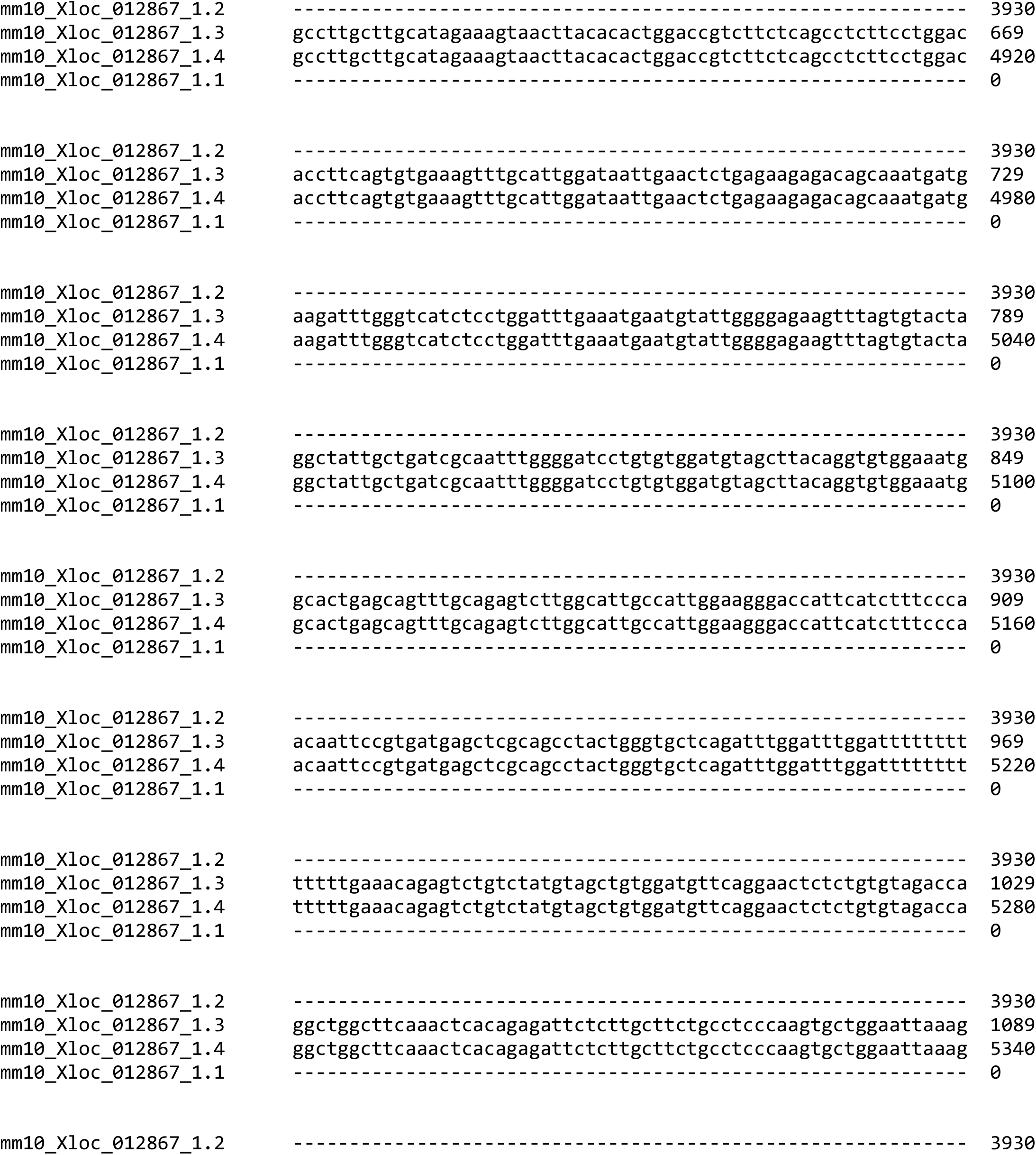
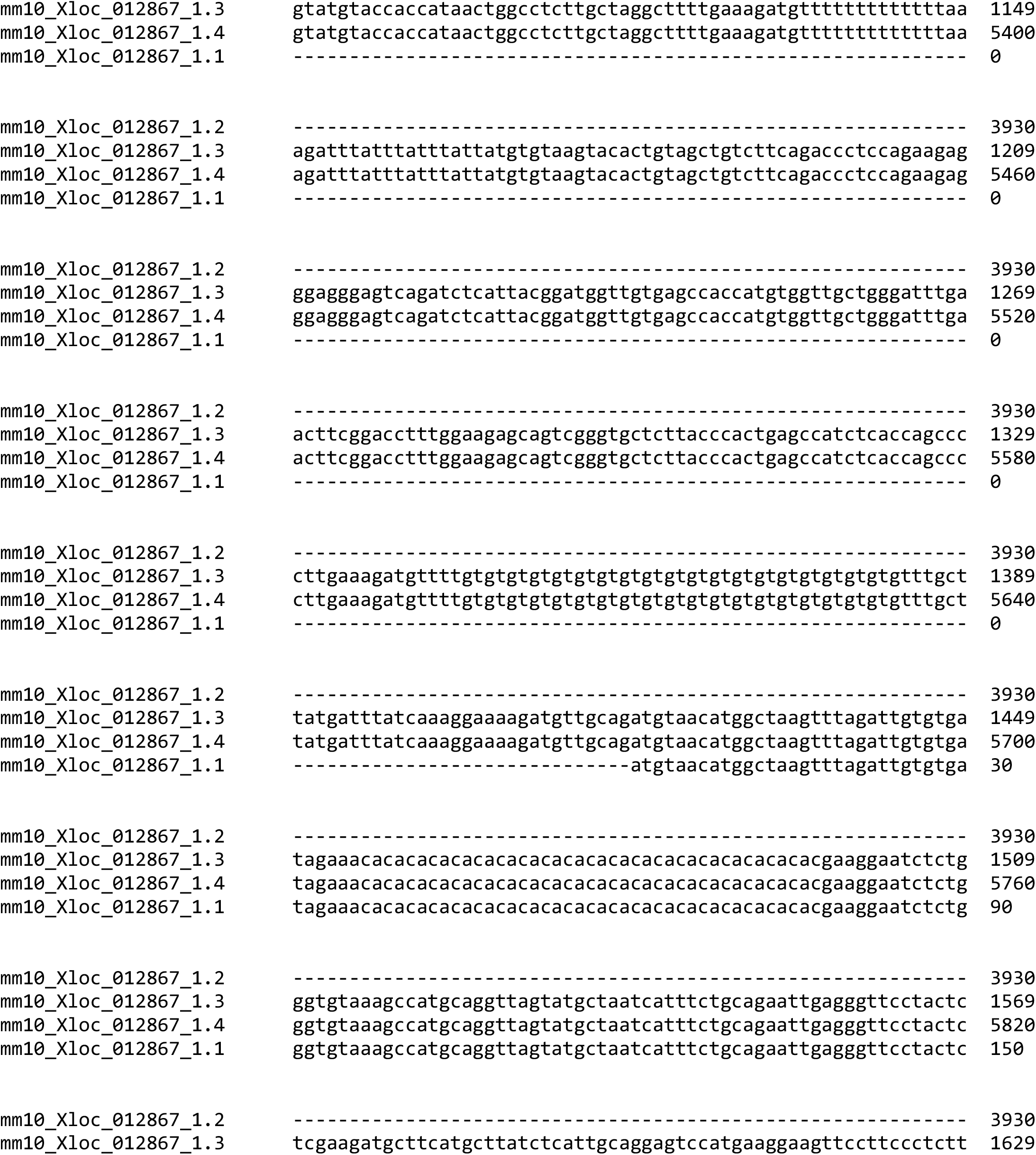
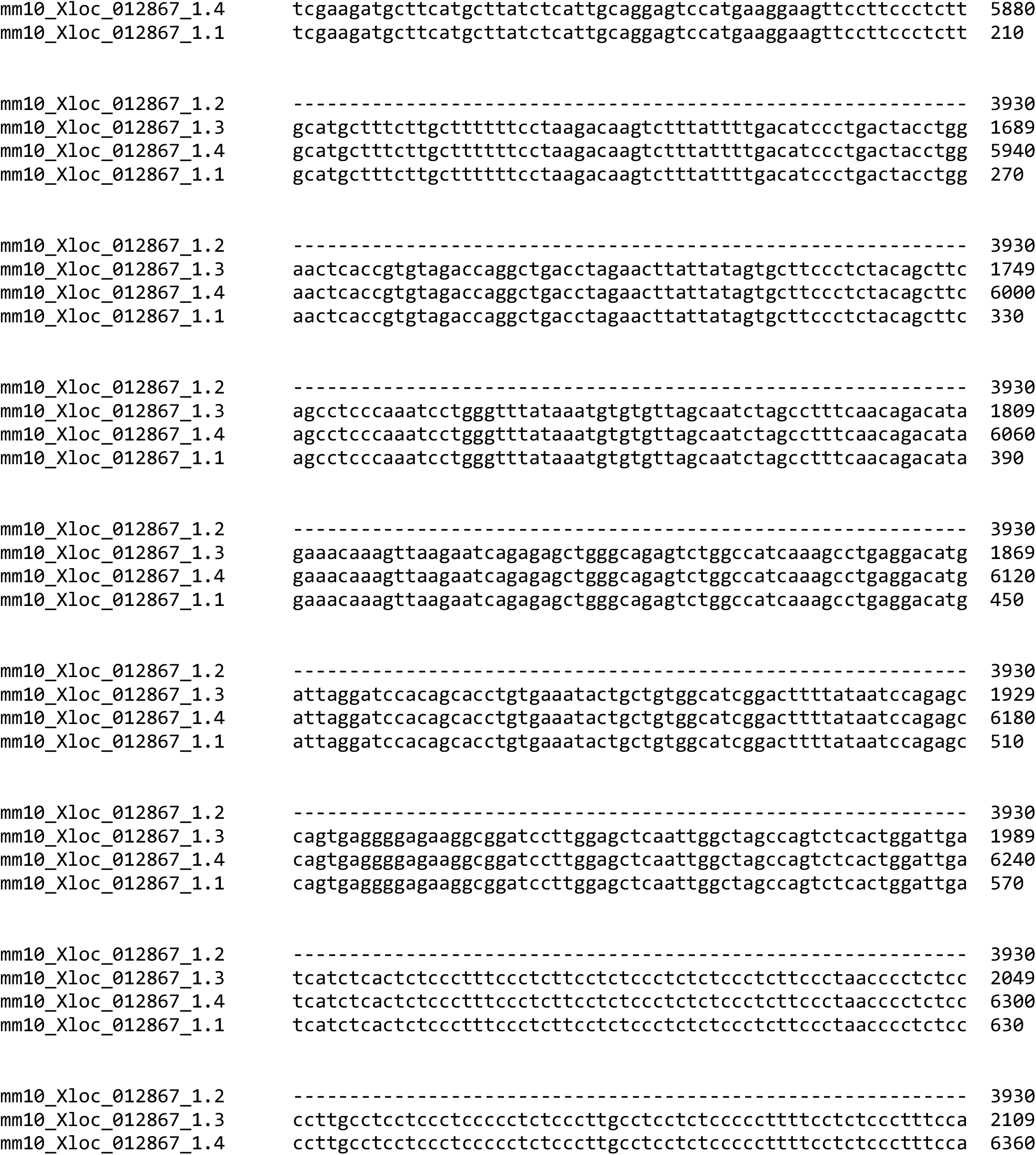
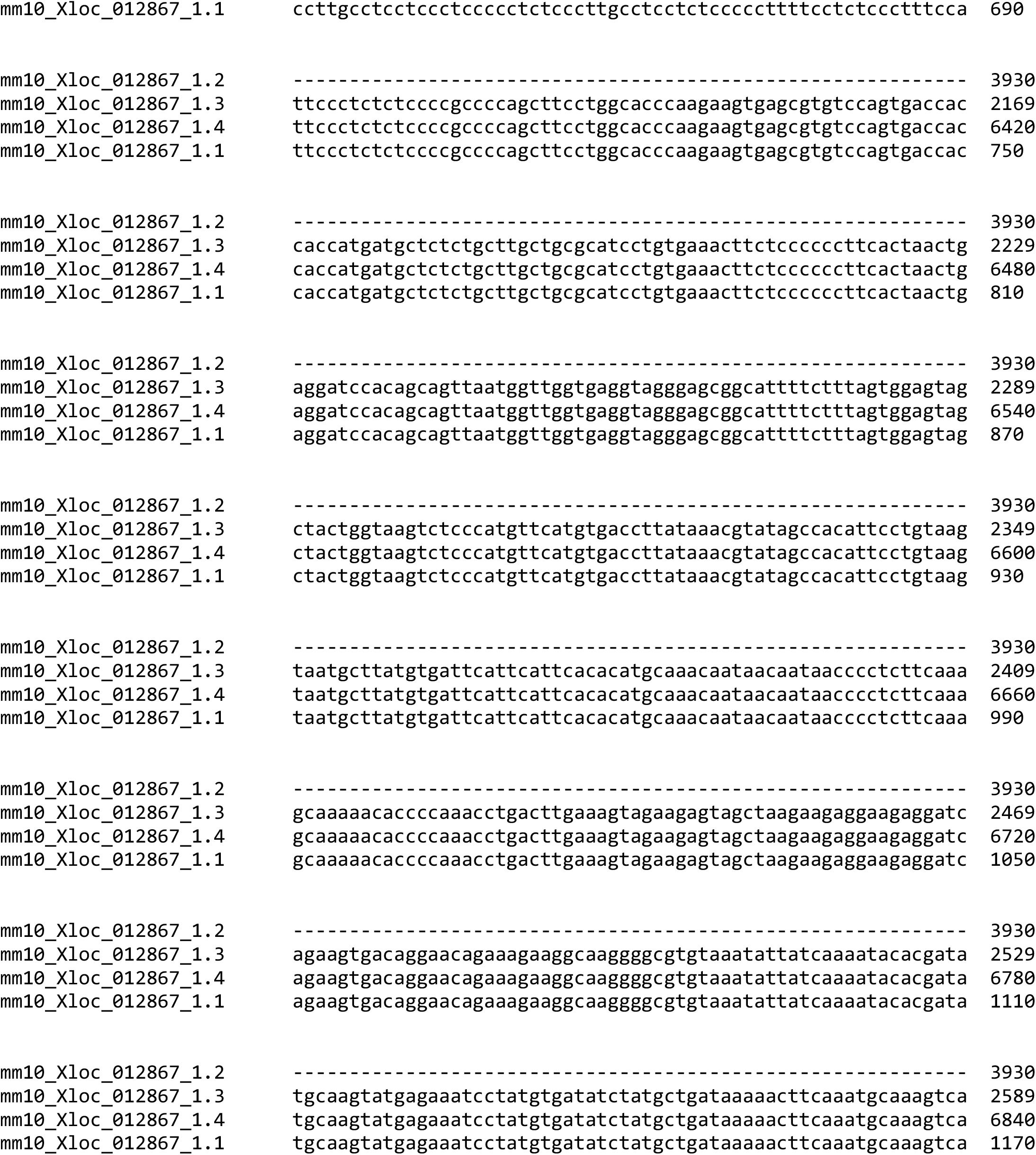
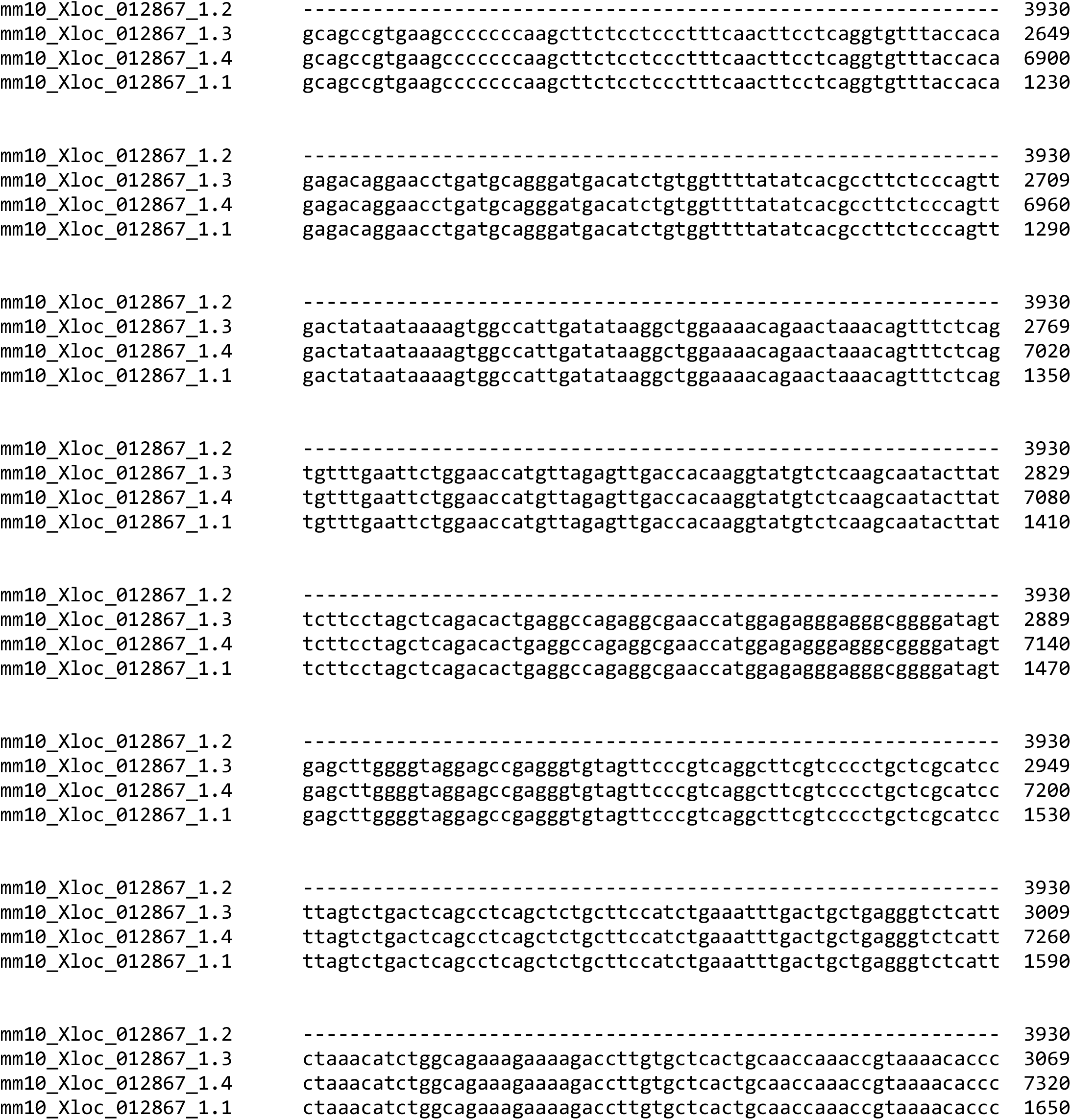
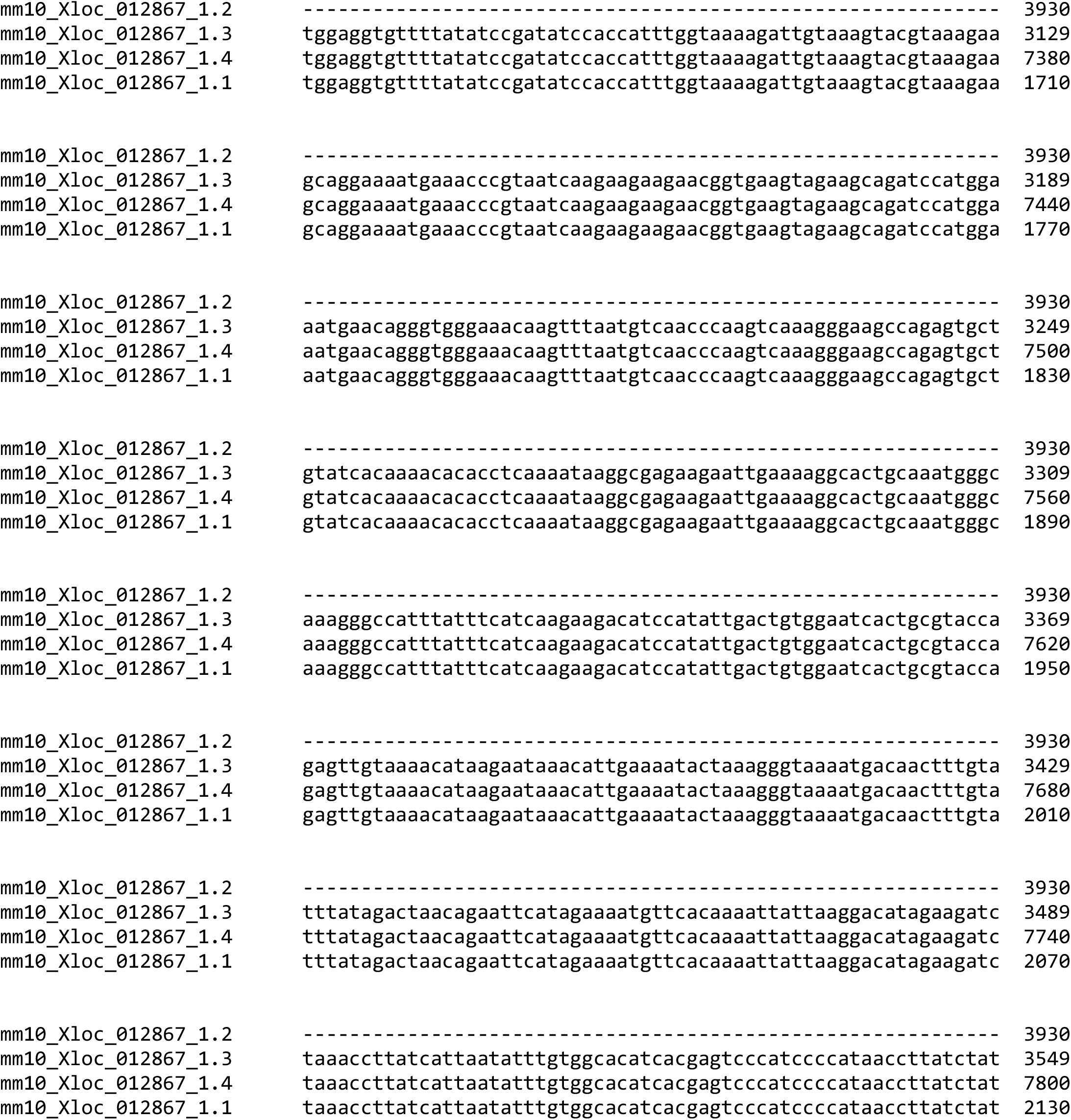
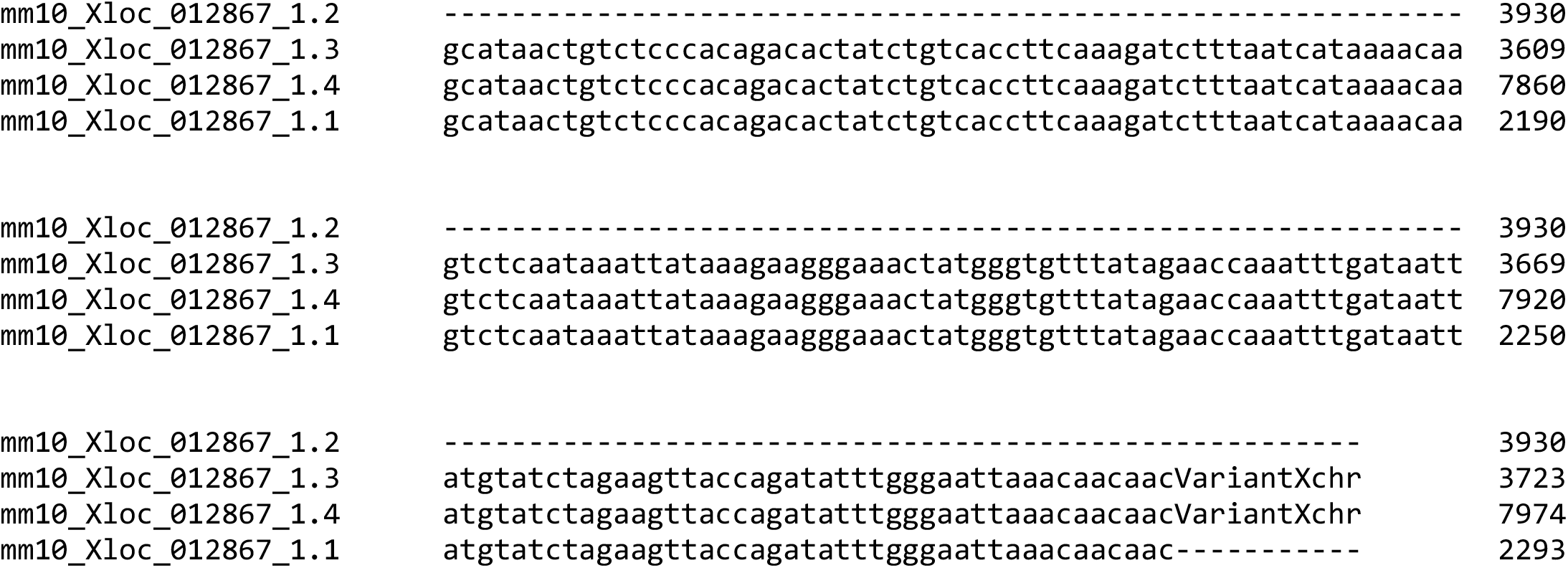

## Additional file 3: Supplementary Information. Methods, Table and Figure.

### Supplementary Methods

miRNA expression analysis was evaluated using TaqMan^TM^ MicroRNA Assay probes (Applied Biosystems). U6 was used as an internal control. *Notch1* primers were designed using the Primer3 online tool (https://www.bioinfo.ut.ee/primer3-0.4.0/) and ordered from IDT. Primer sequences are provided in Table S2 table. Reactions were carried out in the StepOneTM Real-Time PCR System (Thermo Fisher Scientific) and the analysis was performed using StepOneTM software v2.1. At least three independent experiments were conducted with duplicates and a no-template control (NTC) as a negative control. Melt-curve analysis was done in every reaction for the confirmation of a single product. Data was analyzed by the relative quantification (ΔΔCt) method and expressed as mean ± standard error of the mean (SEM).

**Table S2.**
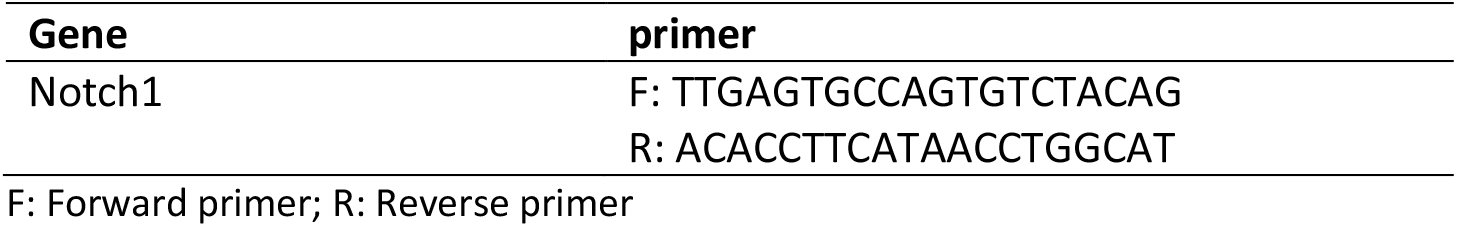
qRT-PCR SYBR® Green primer sequences.

**Fig. S3.**
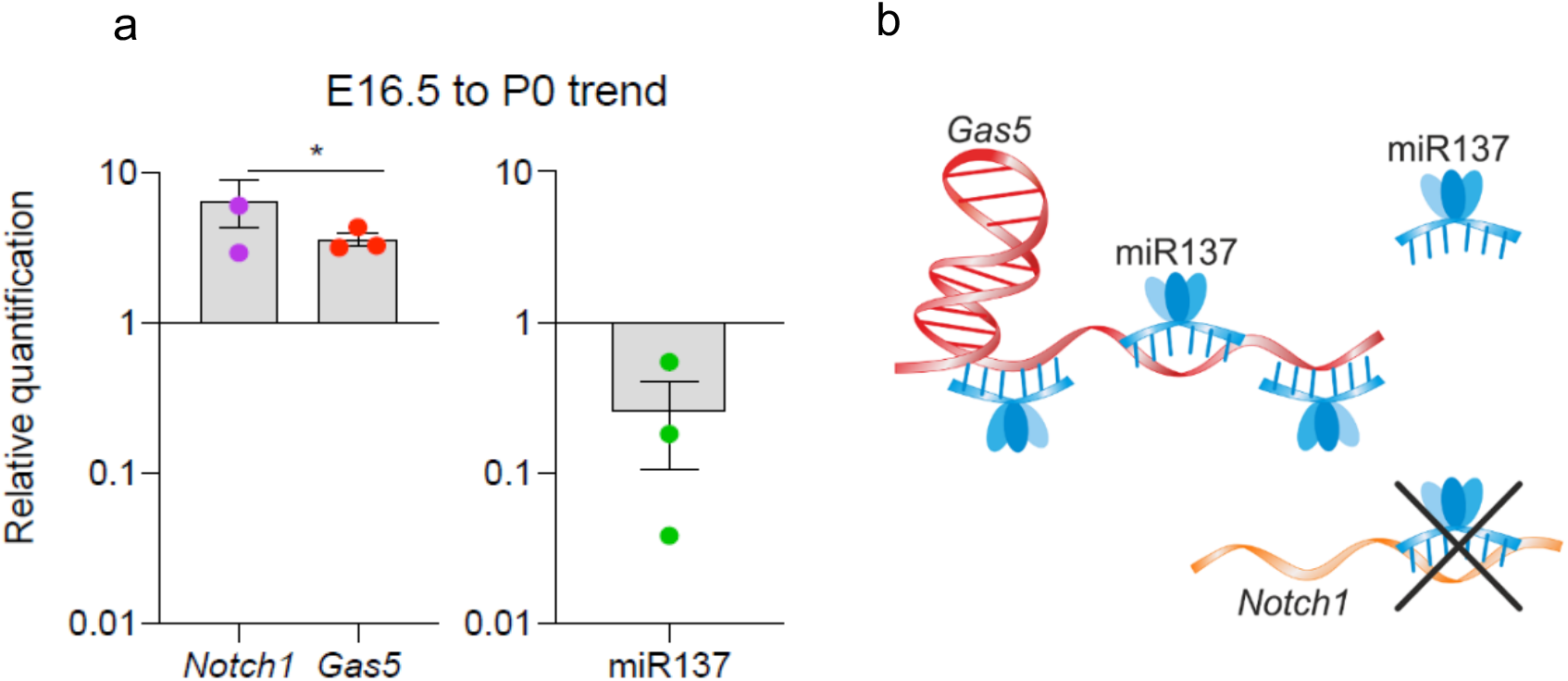
*Gas5* and miR-137 regulation in the Notch pathway. Expression trends between E16.5 and P0 sensory epithelium RNA of **a** *Notch1*, *Gas5*, and miR-137 using qRT-PCR. Data is presented as E16.5 vs. P0 mean ±SEM, n=3. Multiple t-test with Holm-Sidak post-hoc correction for multiple comparisons was applied for statistical analysis. **b** Scheme of predicted pathway.

## Additional file 4: Supplementary Information. Table S3.

**Table S3.**
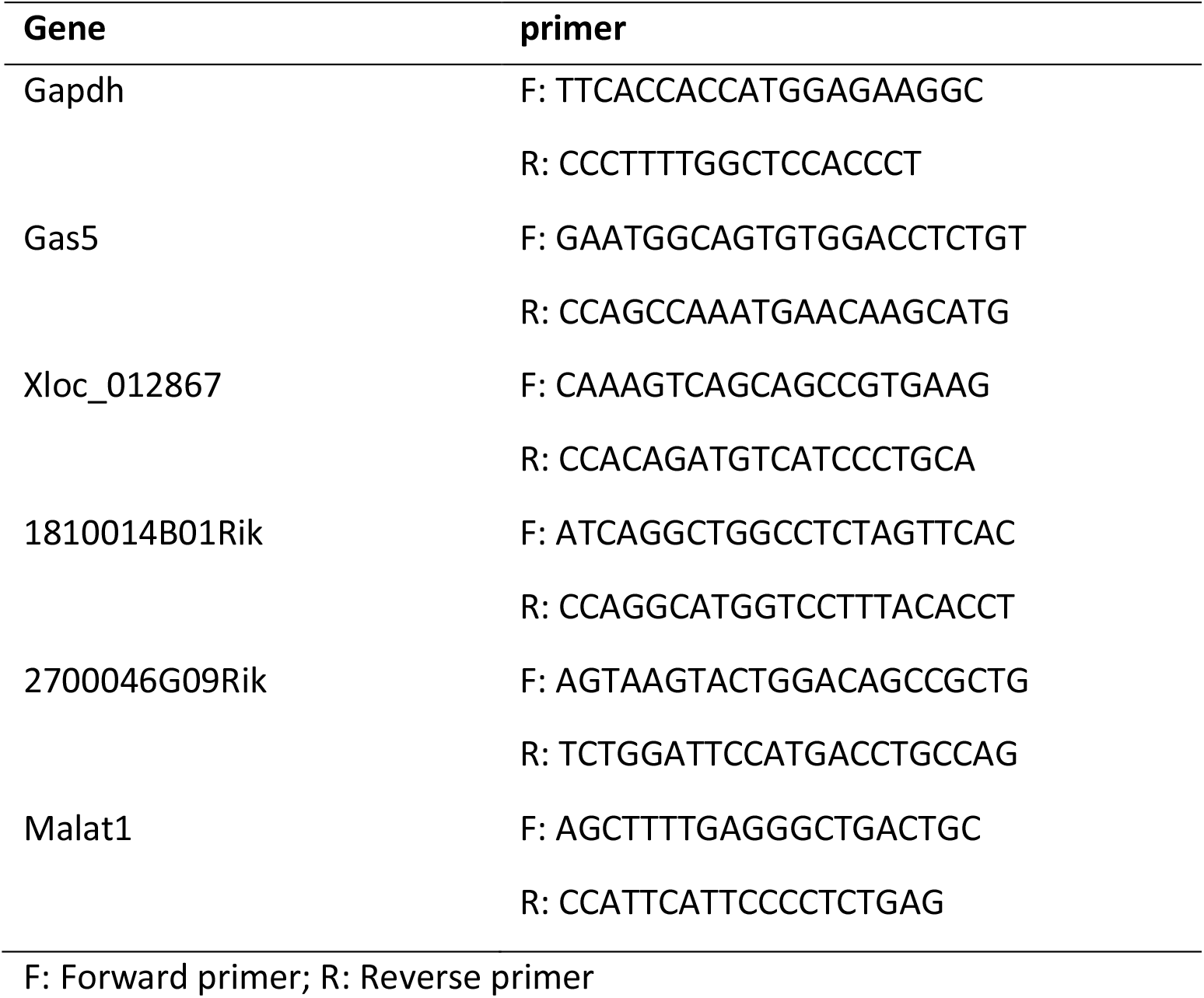
qRT-PCR SYBR® Green primer sequences

## References

[1] Olusanya, BO, Davis, AC, Hoffman HJ. Hearing loss: rising prevalence and impact. Bull World Health Organ. 2019;97:646–A.

[2] Shukla, A, Harper, M, Pedersen, E, et al. Hearing, loss, loneliness, and social isolation: A systematic review. Otolaryngol Head Neck Surg. 2020;194599820910377.

[3] Shearer, AE, Smith RJ. Massively parallel sequencing for genetic diagnosis of hearing loss: The new standard of care. Otolaryngol Head Neck Surg. 2015;153:175–82.

[4] Richardson, GP, de Monvel, JB, Petit C. How the genetics of deafness illuminates auditory physiology. Annu Rev Physiol. 2011;73:311–34.

[5] Naranjo, S, Voesenek, K, de la Calle-Mustienes, E, et al. Multiple enhancers located in a 1-Mb region upstream of *POU3F4* promote expression during inner ear development and may be required for hearing. Hum Genet. 2010;128:411–9.

[6] Mencia, A, Modamio-Hoybjor, S, Redshaw, N, et al. Mutations in the seed region of human miR-96 are responsible for nonsyndromic progressive hearing loss. Nat Genet. 2009;41:609–13.

[7] Schultz, JM, Khan, SN, Ahmed, ZM, et al. Noncoding mutations of *HGF* are associated with nonsyndromic hearing, loss, DFNB39. Am J Hum Genet. 2009;85:25–39.

[8] Guttman, M, Amit, I, Garber, M, et al. Chromatin signature reveals over a thousand highly conserved large non-coding RNAs in mammals. Nature. 2009;458:223–7.

[9] Cabili, MN, Dunagin, MC, McClanahan, PD, et al. Localization and abundance analysis of human lncRNAs at single-cell and single-molecule resolution. Genome Biol. 2015;16:20.

[10] Ulitsky, I, Bartel DP. lincRNAs: Genomics, evolution, and mechanisms. Cell. 2013;154:26–46.

[11] Hu, W, Alvarez-Dominguez, JR, Lodish HF. Regulation of mammalian cell differentiation by long non-coding RNAs. EMBO Rep. 2012;13:971–83.

[12] Kung, JT, Colognori, D, Lee JT. Long noncoding RNAs: past, present, and future. Genetics. 2013;193:651–69.

[13] Beermann, J, Piccoli, MT, Viereck, J, et al. Non-coding RNAs in development and disease: Background, mechanisms, and therapeutic approaches. Physiol Rev. 2016;96:1297–325.

[14] Mustafi, D, Kevany, BM, Bai, X, et al. Evolutionarily conserved long intergenic non-coding RNAs in the eye. Hum Mol Genet. 2013;22:2992–3002.

[15] Young, TL, Matsuda, T, Cepko CL. The noncoding RNA taurine upregulated gene 1 is required for differentiation of the murine retina. Curr Biol. 2005;15:501–12.

[16] Wang, M, Liu, W, Jiao, J, et al. Expression profiling of mRNAs and long non-coding RNAs in aged mouse olfactory bulb. Sci Rep. 2017;7:2079.

[17] Camargo, AP, Nakahara, TS, Firmino, LER, et al. Uncovering the mouse olfactory long non-coding transcriptome with a novel machine-learning model. DNA Res. 2019;26:365–78.

[18] Schrauwen, I, Hasin-Brumshtein, Y, Corneveaux, JJ, et al. A comprehensive catalogue of the coding and non-coding transcripts of the human inner ear. Hear Res. 2016;333:266–74.

[19] Ushakov, K, Koffler-Brill, T, Aviv, R, et al. Genome-wide identification and expression profiling of long non-coding RNAs in auditory and vestibular systems. Sci Rep. 2017;7:8637.

[20] Van Camp, G, Smith RJH. Hereditary Hearing Loss Homepage. http://hereditaryhearinglossorg Accessed June 2020.

[21] Kolla, L, Kelly, MC, Mann, ZF, et al. Characterization of the development of the mouse cochlear epithelium at the single cell level. Nat Commun. 2020;11:2389.

[22] Engreitz, JM, Haines, JE, Perez, EM, et al. Local regulation of gene expression by lncRNA promoters, transcription and splicing. Nature. 2016;539:452–5.

[23] Yizhar-Barnea, O, Valensisi, C, Jayavelu, ND, et al. DNA methylation dynamics during embryonic development and postnatal maturation of the mouse auditory sensory epithelium. Sci Rep. 2018;8:17348.

[24] Lerer, I, Sagi, M, Ben-Neriah, Z, et al. A deletion mutation in *GJB6* cooperating with a *GJB2* mutation in *trans* in non-syndromic deafness: A novel founder mutation in Ashkenazi Jews. Hum Mutat. 2001;18:460.

[25] Estivill, X, Fortina, P, Surrey, S, et al. Connexin-26 mutations in sporadic and inherited sensorineural deafness. Lancet. 1998;351:394–8.

[26] Dinger, ME, Pang, KC, Mercer, TR, et al. Differentiating protein-coding and noncoding RNA: challenges and ambiguities. PLoS Comput Biol. 2008;4:e1000176.

[27] Housman, G, Ulitsky I. Methods for distinguishing between protein-coding and long noncoding RNAs and the elusive biological purpose of translation of long noncoding RNAs. Biochim Biophys Acta. 2016;1859:31–40.

[28] Ramani, R, Krumholz, K, Huang, YF, et al. PhastWeb: a web interface for evolutionary conservation scoring of multiple sequence alignments using phastCons and phyloP. Bioinformatics. 2019;35:2320–2.

[29] Raho, G, Barone, V, Rossi, D, et al. The gas 5 gene shows four alternative splicing patterns without coding for a protein. Gene. 2000;256:13–7.

[30] Mourtada-Maarabouni, M, Pickard, MR, Hedge, VL, et al. GAS5, a non-protein-coding, RNA, controls apoptosis and is downregulated in breast cancer. Oncogene. 2009;28:195–208.

[31] Tu, J, Tian, G, Cheung, HH, et al. Gas5 is an essential lncRNA regulator for self-renewal and pluripotency of mouse embryonic stem cells and induced pluripotent stem cells. Stem Cell Res Ther. 2018;9:71.

[32] Anderson, DM, Anderson, KM, Chang, CL, et al. A micropeptide encoded by a putative long noncoding RNA regulates muscle performance. Cell. 2015;160:595–606.

[33] Lin, MF, Jungreis, I, Kellis M. PhyloCSF: a comparative genomics method to distinguish protein coding and non-coding regions. Bioinformatics. 2011;27:i275–82.

[34] Lin, ST, Huang, Y, Zhang, L, et al. MicroRNA-23a promotes myelination in the central nervous system. Proc Natl Acad Sci USA. 2013;110:17468–73.

[35] Lin, ST, Heng, MY, Ptacek, LJ, et al. Regulation of myelination in the central nervous system by nuclear lamin B1 and non-coding RNAs. Transl Neurodegener. 2014;3:4.

[36] Arisi, I, D’Onofrio, M, Brandi, R, et al. Gene expression biomarkers in the brain of a mouse model for Alzheimer’s disease: mining of microarray data by logic classification and feature selection. J Alzheimers Dis. 2011;24:721–38.

[37] Wang, F, Flanagan, J, Su, N, et al. RNAscope: a novel in situ RNA analysis platform for formalin-fixed, paraffin-embedded tissues. J Mol Diagn. 2012;14:22–9.

[38] Groves, AK, Fekete DM. Shaping sound in space: the regulation of inner ear patterning. Development. 2012;139:245–57.

[39] Consortium EP. A user’s guide to the encyclopedia of DNA elements (ENCODE). PLoS Biol. 2011;9:e1001046.

[40] Baer, C, Claus, R, Plass C. Genome-wide epigenetic regulation of miRNAs in cancer. Cancer Res. 2013;73:473–7.

[41] Castellanos-Rubio, A, Fernandez-Jimenez, N, Kratchmarov, R, et al. A long noncoding RNA associated with susceptibility to celiac disease. Science. 2016;352:91–5.

[42] Gomes, CPC, Schroen, B, Kuster, GM, et al. Regulatory RNAs in heart failure. Circulation. 2020;141:313–28.

[43] Kopp, F, Mendell JT. Functional classification and experimental dissection of long noncoding RNAs. Cell. 2018;172:393–407.

[44] Ip, JY, Nakagawa S. Long non-coding RNAs in nuclear bodies. Dev Growth Differ. 2012;54:44–54.

[45] Iyer, MK, Niknafs, YS, Malik, R, et al. The landscape of long noncoding RNAs in the human transcriptome. Nat Genet. 2015;47:199–208.

[46] Sarropoulos, I, Marin, R, Cardoso-Moreira, M, et al. Developmental dynamics of lncRNAs across mammalian organs and species. Nature. 2019;571:510–4.

[47] Manji, SS, Sorensen, BS, Klockars, T, et al. Molecular characterization and expression of maternally expressed gene 3 (*Meg3/Gtl2*) RNA in the mouse inner ear. J Neurosci Res. 2006;83:181–90.

[48] Roberts, KA, Abraira, VE, Tucker, AF, et al. Mutation of *Rubie*, a novel long non-coding RNA located upstream of *Bmp4*, causes vestibular malformation in mice. PLoS One. 2012;7:e29495.

[49] Wesdorp, M, de Koning Gans, PAM, Schraders M, et al. Heterozygous missense variants of *LMX1A* lead to nonsyndromic hearing impairment and vestibular dysfunction. Hum Genet. 2018;137:389–400.

[50] Zhang, S, Reljic, B, Liang, C, et al. Mitochondrial peptide BRAWNIN is essential for vertebrate respiratory complex III assembly. Nat Commun. 2020;11:1312.

[51] Cesana, M, Cacchiarelli, D, Legnini, I, et al. A long noncoding RNA controls muscle differentiation by functioning as a competing endogenous RNA. Cell. 2011;147:358–69.

[52] Rinn, JL, Chang HY. Genome regulation by long noncoding RNAs. Annu Rev Biochem. 2012;81:145–66.

[53] Kino, T, Hurt, DE, Ichijo, T, et al. Noncoding RNA gas5 is a growth arrest- and starvation-associated repressor of the glucocorticoid receptor. Sci Signal. 2010;3:ra8.

[54] Kelsell, DP, Dunlop, J, Stevens, HP, et al. Connexin 26 mutations in hereditary non-syndromic sensorineural deafness. Nature. 1997;387:80–3.

[55] Wienholds, E, Kloosterman, WP, Miska, E, et al. MicroRNA expression in zebrafish embryonic development. Science. 2005;309:310–1.

[56] Xu, S, Witmer, PD, Lumayag, S, et al. MicroRNA (miRNA) transcriptome of mouse retina and identification of a sensory organ-specific miRNA cluster. J Biol Chem. 2007;282:25053–66.

[57] Weston, MD, Pierce, ML, Jensen-Smith, HC, et al. MicroRNA-183 family expression in hair cell development and requirement of microRNAs for hair cell maintenance and survival. Dev Dyn. 2011;240:808–19.

[58] Chen, F, Zhang, L, Wang, E, et al. LncRNA GAS5 regulates ischemic stroke as a competing endogenous RNA for miR-137 to regulate the Notch1 signaling pathway. Biochem Biophys Res Commun. 2018;496:184–90.

[59] Shu, Y, Li, W, Huang, M, et al. Renewed proliferation in adult mouse cochlea and regeneration of hair cells. Nat Commun. 2019;10:5530.

[60] Brown, R, Groves AK. Hear, hear for notch: Control of cell fates in the inner ear by notch signaling. Biomolecules. 2020;10:

[61] Kelley MW. Cellular commitment and differentiation in the organ of Corti. Int J Dev Biol. 2007;51:571–83.

1. World Health Organization, http://www.who.int/news-room/fact-sheets/detail/deafness-and-hearing-loss. Accessed July 2020.

## References

1. Tani H, Torimura M, Akimitsu, N. The RNA degradation pathway regulates the function of GAS5 a non-coding RNA in mammalian cells. PLoS One. 2013;8:e55684.

2. Ulitsky I, Shkumatava A, Jan CH, Sive H, Bartel, DP. Conserved function of lincRNAs in vertebrate embryonic development despite rapid sequence evolution. Cell. 2011;147:1537–50.

3. Ruan W, Wang P, Feng S, Xue Y, Li, Y. Long non-coding RNA small nucleolar RNA host gene 12 (SNHG12) promotes cell proliferation and migration by upregulating angiomotin gene expression in human osteosarcoma cells. Tumour Biol. 2016;37:4065–73.

4. Acevedo-Alvarez M, Yeh J, Alvarez-Lugo L, Lu M, Sukumar N, Hill WG et al. Mouse urothelial genes associated with voiding behavior changes after ovariectomy and bladder lipopolysaccharide exposure. Neurourol Urodyn. 2018;37:2398–405.

5. Zhang D, Cao C, Liu L, Wu, D. Up-regulation of lncRNA SNHG20 predicts poor prognosis in hepatocellular carcinoma. J Cancer. 2016;7:608–17.

6. Fan J, Zhou Q, Li Y, Song X, Hu J, Qin Z et al. Profiling of Long non-coding RNAs and mRNAs by RNA-sequencing in the hippocampi of adult mice following propofol sedation. Front Mol Neurosci. 2018;11:91.

7. Arisi I, D’Onofrio M, Brandi R, Felsani A, Capsoni S, Drovandi G et al. Gene expression biomarkers in the brain of a mouse model for Alzheimer’s disease: mining of microarray data by logic classification and feature selection. J Alzheimers Dis. 2011;24:721–38.

8. Lin ST, Huang Y, Zhang L, Heng MY, Ptacek LJ, Fu, YH. MicroRNA-23a promotes myelination in the central nervous system. Proc Natl Acad Sci USA. 2013;110:17468–73.

9. Paralkar VR, Taborda CC, Huang P, Yao Y, Kossenkov AV, Prasad R et al. Unlinking an lncRNA from its associated cis element. Mol Cell. 2016;62:104–10.

10. Zhang M, Wang W, Li T, Yu X, Zhu Y, Ding F et al. Long noncoding RNA SNHG1 predicts a poor prognosis and promotes hepatocellular carcinoma tumorigenesis. Biomed Pharmacother. 2016;80:73–9.

11. Yi H, Peng R, Zhang LY, Sun Y, Peng HM, Liu HD et al. LincRNA-Gm4419 knockdown ameliorates NF-kappaB/NLRP3 inflammasome-mediated inflammation in diabetic nephropathy. Cell Death Dis. 2017;8:e2583.

